# Spatial clustering of inhibition in mouse primary visual cortex

**DOI:** 10.1101/608125

**Authors:** Pawan Bista, Rinaldo D. D’Souza, Andrew M. Meier, Weiqing Ji, Andreas Burkhalter

**Author notes:** Correspondence: Andreas Burkhalter PhD, Department of Neuroscience, 8108, Washington University School of Medicine, 660 S. Euclid Avenue, St. Louis, MO 63110. Co-first author (Pawan Bista, Rinaldo D. D’Souza).

## Abstract

Whether mouse visual cortex contains orderly feature maps is debated. The overlapping pattern of geniculocortical (dLGN) inputs with M2 muscarinic acetylcholine receptor-rich patches in layer 1 (L1) suggests a non-random architecture. Here, we found that L1 inputs from the lateral posterior thalamus (LP) avoid patches and target interpatches. Channelrhodopsin-assisted mapping of EPSCs in L2/3 shows that the relative excitation of parvalbumin-expressing interneurons (PVs) and pyramidal neurons (PNs) by dLGN, LP and cortical feedback are distinct and depend on whether the neurons reside in clusters aligned with patches or interpatches. Paired recordings from PVs and PNs shows that unitary IPSCs are larger in interpatches than patches. The spatial clustering of inhibition is matched by dense clustering of PV-terminals in interpatches. The results show that the excitation/inhibition balance across V1 is organized into patch and interpatch subnetworks which receive distinct long-range inputs and are specialized for the processing of distinct spatiotemporal features.

## INTRODUCTION

The modernist’s maxim that form follows function manifests itself in neuroscience as functional architecture. The mesoscale description of the spatial relationship between neuronal responses, layers, columns and areas, has driven much of what is known about the matrix of the cortical machinery (Hubel and Wiesel, 1962, 1974). The concepts on the network organization that emerged from these foundational studies include: the orderly representation of space in topographic maps, the complexification of receptive fields across layers, functionally distinct columns, distributed hierarchical processing along specialized functional streams, the exponential distance rule of cortical connectivity, and the selection of sensory input based on feedback from higher areas (Felleman and Van Essen, 1991; Horvát et al., 2016; Hubel and Wiesel, 1962). Although the functional architecture implies that the spatiotemporal tuning of cell populations is determined by their connectivity and physical layout, the underlying networks of modular processing have been difficult to define (DaCosta and Martin, 2013). The challenge has been greatest in rodents because of the long-held view that cortex lacks columnar organization (Ohki et al., 2005; 2007). However, recent studies in mouse primary visual cortex (V1) have found that subcortically and intracortically projecting PNs are spatially clustered and are vertically aligned to mini- and microcolumns of cell bodies and dendrites with distinct tuning preferences (Kondo et al, 2016; Marukoda et al., 2017; Znamenskiy et al. 2018).

Previously, we found a fixed interdigitating pattern of M2 muscarinic acetylcholine receptor-expressing (M2+) patches and M2- interpatches in L1 of mouse V1, which aligns with functionally distinct cell clusters in L2/3 tuned for high spatial frequency (SF) and high temporal frequency (TF), respectively (Ji et al., 2015). This finding provides structural evidence for functionally discrete modules and raises the question whether the excitation (E) / inhibition (I) balance in patches and interpatches with diverse spatiotemporal stimulus preferences is spatially organized. That inhibition is not a uniform blanket across V1 (Karnani et al., 2014), but is deployed in spatially clustered patterns by subtypes of PV or somatostatin-(SOM) expressing GABAergic neurons (Ebina et al., 2014; Maruoka et al., 2017), and that activation of these cells including those which express vasoactive intestinal peptide can shape stimulus selectivities of PNs is gradually gaining acceptance (Ayzenshtat et al., 2016; Lee, et al., 2012; Lee et al. 2014; Zhu et al., 2015). Whether the inhibitory network is tied to the spatially clustered patch/interpatch system in V1 and provides distinct subnetworks for processing visual information remains unknown.

To determine whether inhibition across V1 is modular we measured the strength of synaptic long-range input to L2/3 PNs and PVs in patches and interpatches by using subcellular Channelrhodopsin-2 (ChR2) assisted circuit mapping (sCRACM) (Mao et al, 2011; Petreanu et al., 2009). We recorded from pairs of PVs and PNs and analyzed unitary excitatory (uEPSCs) and inhibitory (uIPSCs) postsynaptic currents in patches and interpatches. The results show that V1 contains two different circuit motifs in which patches and interpatches have distinct thalamocortical and interareal inputs to PNs and PVs, and that PVs in interpatches provide stronger feedforward inhibition (FFI) to PNs than in patches. The modular organization of inhibition is consistent with the notion that neurons in interpatches are more sensitive to rapidly changing visual inputs (Ji et al., 2015).

## RESULTS

### Clustering of thalamocortical inputs to L1 of V1

We have recently shown that inputs from the dLGN to L1 of V1 are spatially clustered and overlap with the patchy pattern of M2 immunostaining (Ji et al., 2015). Here, we show in tangential sections of flatmounted cortex of Chrm2tdT mice that dLGN→V1 axons labeled by anterograde tracing with AAV2/1.hSyn.EGFP preferentially terminate in M2+ patches of L1 and avoid M2- interpatches (Figure 1A-C). For quantitative analysis we divided images into patches and interpatches based on M2 fluorescence intensity contour maps (Sincich and Horton, 2005) (Figure S1), and measured EGFP intensity in patches (top 3^rd^) and interpatches (bottom 3^rd^). We then normalized the pixel values in interpatches to the mean intensity in patches and plotted the counts in different intensity bins. We found that the fluorescence intensity in patches was 2.1 ± 0.024-fold (*p* = 8×10^−18^, Komolgorov-Smirnov test [KS]) higher in patches than interpatches (Figure 1D). Similar to our previous findings, patches and interpatches in L1 were 60-80 μm wide and their centroids were 120-140 μm apart. dLGN projections to L4, and 5/6 appeared uniform (data not shown). Inputs to L1 from the LP thalamus exhibited a strikingly different pattern, showing 1.6 ± 0.14-fold (*p* = 1.33×10^−4^, KS) stronger projections to M2- interpatches (Figure 1E-H). Simultaneous tracing of dLGN and LP inputs with AAV2/1hSyn.tdTomato.WPRE.bGH and AAV2/1.hSyn.EGFP, respectively, confirmed the interdigitating pattern of projections from primary and higher order thalamic nuclei, showing denser LP input to interpatches (*p* = 7.95 x 10^−4^, KS) (Figure 1I-M).

**Figure 1.**
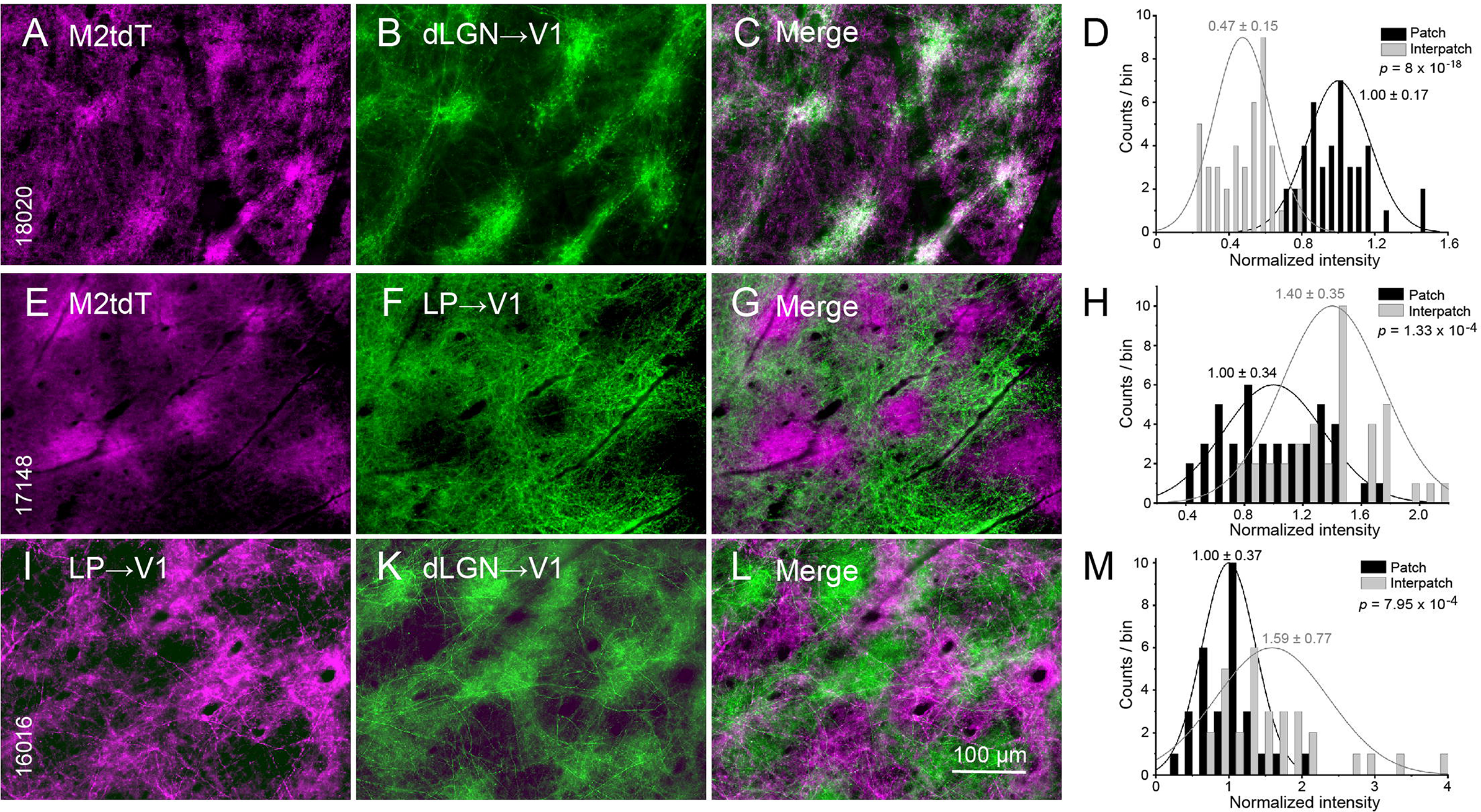
Anterograde viral tracing of dLGN and LP projections to L1 of V1. (A-C, E-G, I-L) Tangential sections through L1 showing M2+ patches and M2-interpatches in V1 of Chrm2tdT (A-C, E-G) and C57BL/6J mice (I-L). (A-C) Clustered dLGN projections traced with AAV2/1.hSyn.EGFP (green) overlap with M2tdT patches (purple). (D) Fluorescence intensity of dLGN input to patches (normalized to patches) is higher than to interpatches. (E-G) Clustered LP projections traced with AAV2/1.hSyn.EGFP (green) overlap with M2- interpatches. (H) Fluorescence intensity of LP inputs to interpatches (normalized to patches) is higher than to patches. (I-L) Interdigitating projections from LP to interpatches traced with AAV2/1hSyn.tdTomato.WPRE.bGH (purple) and dLGN traced with AAV2/1.hSyn.EGFP (green) to patches. (M) Dense LP projections to interpatches where dLGN input is weak. Mean ± SD, KS test (D, H, M).

### Clustering of intracortical inputs to L1 of V1

We next compared feedback projections from the higher visual cortical ventral stream lateromedial area, LM, to V1 with inputs from the dLGN. Double viral tracings from the dLGN (AAV2/1.hSyn.EGFP) and LM (AAV2/1hSyn.tdTomato.WPRE.bGH) showed that inputs from both sources overlapped in presumtive M2+ patches of L1 (Figure 1A-C; 2A). Quantitative analysis showed that LM inputs to patches were 1.7 ± 0.05-fold denser than to interpatches (*p* = 1.45×10^4^, KS) (Figure 2B). We have shown previously that inputs from the dorsal stream anterolateral area, AL, terminate in M2+ patches (Ji et al., 2015), raising the question whether M2+ patches are the preferred targets of cortical feedback. To address this, we traced the connections to V1 from the posteromedial area, PM, another member of the dorsal stream (Wang et al. 2012). We found that inputs from PM were non-uniform, overlapped in part with patches and interpatches but avoided the top and bottom intensity quantile of each compartment (Figure 2F-G). The spatial distribution of PM→V1 inputs showed no difference between patches and interpatches (Figure 2H). However, simultaneous labeling of inputs from AL and PM showed an interdigitating pattern and revealed a fine-scale pattern in patches (Figure 2 I-L). This demonstrates that projections from AL targets the top and PM the bottom intensity quantile of patches and indicates that the terminal patterns of feedback to V1 by the two dorsal stream areas is not identical.

**Figure 2.**
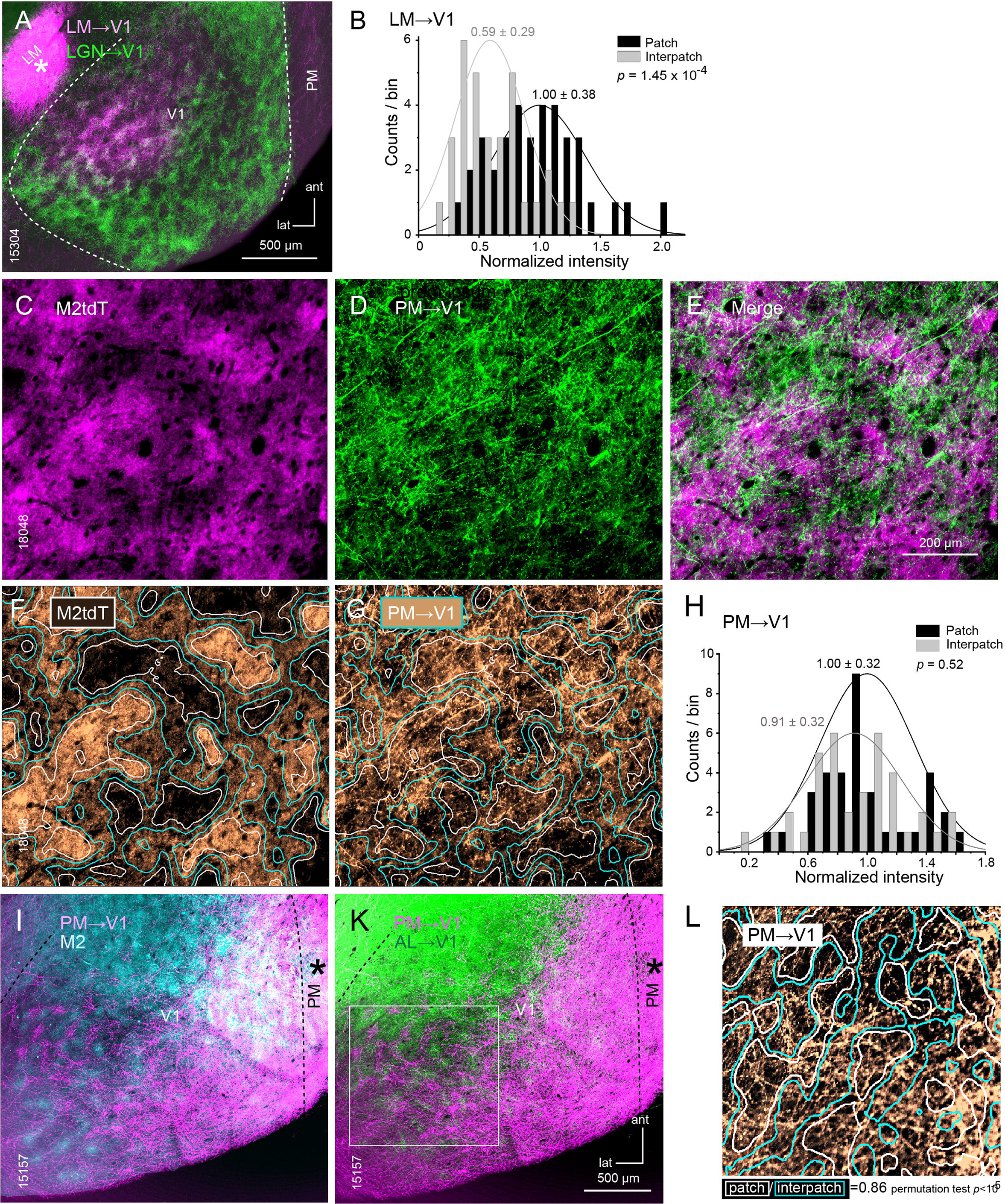
Anterograde viral tracing of LM and PM projections to L1 of V1. (A, C-E, F, G) Tangential sections through L1 of V1 of C57BL/6J (A, I, K) and Chrm2tdT mice (C-E). (A) Overlapping clusters of dLGN projections traced with AAV2/1.hSyn.EGFP (green) and LM input traced by injection [*] of AAV2/1hSyn.tdTomato.WPRE.bGH (purple) to L1 of V1. (B) Fluorescence intensity of LM inputs to interpatches (normalized to patches) is weaker than to patches (M2+, Figure 1A-C). (C-E) Nonuniform distribution of inputs from PM traced with AAV2/1.hSyn.EGFP (green) partially overlapping with M2+ patches (purple) and M2- interpatches. (F, G) Contour maps of M2tdT intensity. White boundaries indicate top (peaks of patches) and bottom (peaks of interpatches) quantiles. Cyan boundaries outline the middle two quantiles. (G) PM inputs to V1 (gold) fall preferentially into zone between top and bottom quantiles. (H) Intensity profile shows that PM inputs to V1 overlap with patches and interpatches. Mean ± SD. KS test. (I, K) Simultaneous tracing of PM (traced with AAV2/1hSyn.tdTomato.WPRE.bGH, purple) and AL (traced with AAV2/1.hSyn.EGFP, green) inputs to V1 superimposed onto immunolabeling for M2 (blue). At low magnification AL and PM projections show an interdigitating pattern. (L) Analysis of projection densities (boxed area in K) at high magnification shows that PM inputs (gold) involve patches (white outlines top 2 quantiles) and interpatches (cyan outlines bottom 2 quantiles). While AL projections avoid interpatches (Ji et al., 2015), note that AL and PM projections to patches show an interdigitating pattern.

### Clustering of cholinergic inputs to L1 of V1

Classic studies suggest that the cholinergic innervation of cortex is diffuse, signaling is through volume transmission, and that spatial selectivity arises from spatial clustering of receptors and axons (Muñoz and Rudy, 2014). But whether cholinergic fibers in cortex are spatially mapped in systematic fashion is unknown. To address the question we stained tangential sections through V1 with antibodies against M2 and choline acetyl transferase ChAT. We found that ChAT labeled axons in L1 were clustered and significantly (*p* = 0.009) denser in M2+ patches (Figure S2A-D).

### Development of M2 clusters

Motivated by the report of Maruoka et al., (2017) that L5 neurons in V1 of 6-day-old (P6) mice form 20 μm-wide microcolumns, we have looked for clustered M2 expression in early postnatal development. Our results in tangential sections through V1 of P4 Chrm2tdT mice show that M2 expression in L1 is patchy (10 −30 μm wide, 52-79 μm center-to-center, Figure S3A). In L2/3 patches were smaller and contained multiple 0.8-1.5 μm wide rings, presumably representing membranes of cross-sectioned ascending dendrites (Figure S3B) (Kondo et al., 2016). At P10 the L1 patches were larger (30 - 60 μm wide) and more widely spaced (80-100 μm center-to-center) (Figure S3C), a difference which may be accounted for by the 1.7 fold increase in brain size (Agrawal et al., 1968).

### Module- and pathway-specific synaptic strengths of inputs to PNs and PVs

We have shown previously that L2/3 neurons in interpatches of V1 are more often tuned to the direction of visual motion and respond to higher speeds and TFs than neurons in patches (Ji et al., 2015). This non-uniform distribution of selectivities suggested that the two cell clusters differentially process spatiotemporal information. FFI mediated by rapid synaptic activation of PVs (D’Souza et al., 2016) is a circuit motif shown to shape the temporal sensitivity of PNs in auditory cortex independent of stimulus adaptation (Li et al., 2014; Natan et al., 2017). Because the strength of FFI depends on the excitatory input to PNs and PVs and the latter’s inhibition of PNs (Atallah et al., 2012), we compared the strength of excitatory inputs to patches and interpatches from different thalamocortical and intracortical pathways to L2/3 neurons in V1. To do this we used sCRACM in acute slices of PVtdT mice in which inputs to V1 were labeled by anterograde tracing with AAV2/1.hSyn.ChR2(H134R).eYFP.WPRE.hGH.

#### dLGN input in tangential slices

To select for inputs to distal dendrites of L2/3 neurons in L1-2 and optimally preserve their 3D organization, we obtained tangential slices of V1. Whole cell patch clamp recordings were performed from pairs of PNs and PVs (< 50 μm apart) aligned with densely ChR2-eYFP-expressing patches or sparsely labeled interpatches, visualized by anterograde viral tracing from the dLGN (Figure 1A). The locations of the recorded neurons were identified by intracellular filling with Alexa-594 hydrazide and *post hoc* immunostaining of re-sectioned slices with an antibody against M2 (Figures 3A, S4A-C). Input strength was measured by EPSCs elicited by laser stimulation of ChR2-expressing axon terminals. Blue light flashes were delivered in an 8×8 grid with 75 μm spacing centered on the cell body. sCRACM mapping in the same slice of dLGN inputs to neighboring PVs and PNs showed stronger synaptic activation in patches than interpatches (Figures 3A, B, D, G), which is consistent with the dense dLGN projections to patches (Figure 1B). Direct comparison of cell pairs in patches and interpatches showed that EPSCs from PVs were larger than from PNs (Figures 3B, D). The population difference is shown in plots in which responses of cell pairs are represented by single points (Figure 3C, E). The geometric mean of EPSCPN/EPSCPV ratios in patches and interpatches is indicated by a red line intersecting the origin of the graph. Input resistance-corrected data is shown by a blue line. The average EPSC recorded from PVs (323.6±130 pA) in patches was 2.3-fold larger (*p* < 0.001, n = 17 pairs, t-test) than from PNs (140.6±81.6 pA). A similar 2.3-fold difference (*p* < 0.001, n = 17 pairs, t-test) was found in interpatches between the more weakly responsive PVs (147.7±91.2 pA) and PNs (62.9±31.1 pA). Heatmaps of EPSCs in patches and interpatches show that responses were maximal near the cell body and decreased toward the distal tip of dendrites (Figures 3B, D). The responsive area was significantly smaller for PNs than PVs (paired t-test, *p* < 0.001), but similar in patches and interpatches (Figure 3F). dLGN inputs to PVs evoked higher current densities (*p* < 0.01, paired t-test) and significantly (*p* < 0.05, paired t-test) faster rise times of EPSCs than inputs to PNs (Figures 3F, S5A). These results show that in tangential slices the strength of synaptic activation by dLGN inputs depends on cell type, the modularity of innervation and the strong preference for module-specific branching of apical dendrites in L1-2 (Figure 1A-D, 6 B-N, S4D, E, I, J). Indeed the combined length of PN (n = 12) and PV (n = 12) dendrites in patches was 3.4- fold higher (*p* < 0.001, t-test) in patches (812 ± 44 μm) than in interpatches (239 ± 29 μm). For cells in interpatches (n =1) the dendritic length in interpatches (898 ± 41 μm) was 9.8-fold higher (*p* < 0.0001, t-test) than in patches (91 ± 32 μm).

**Figure 3.**
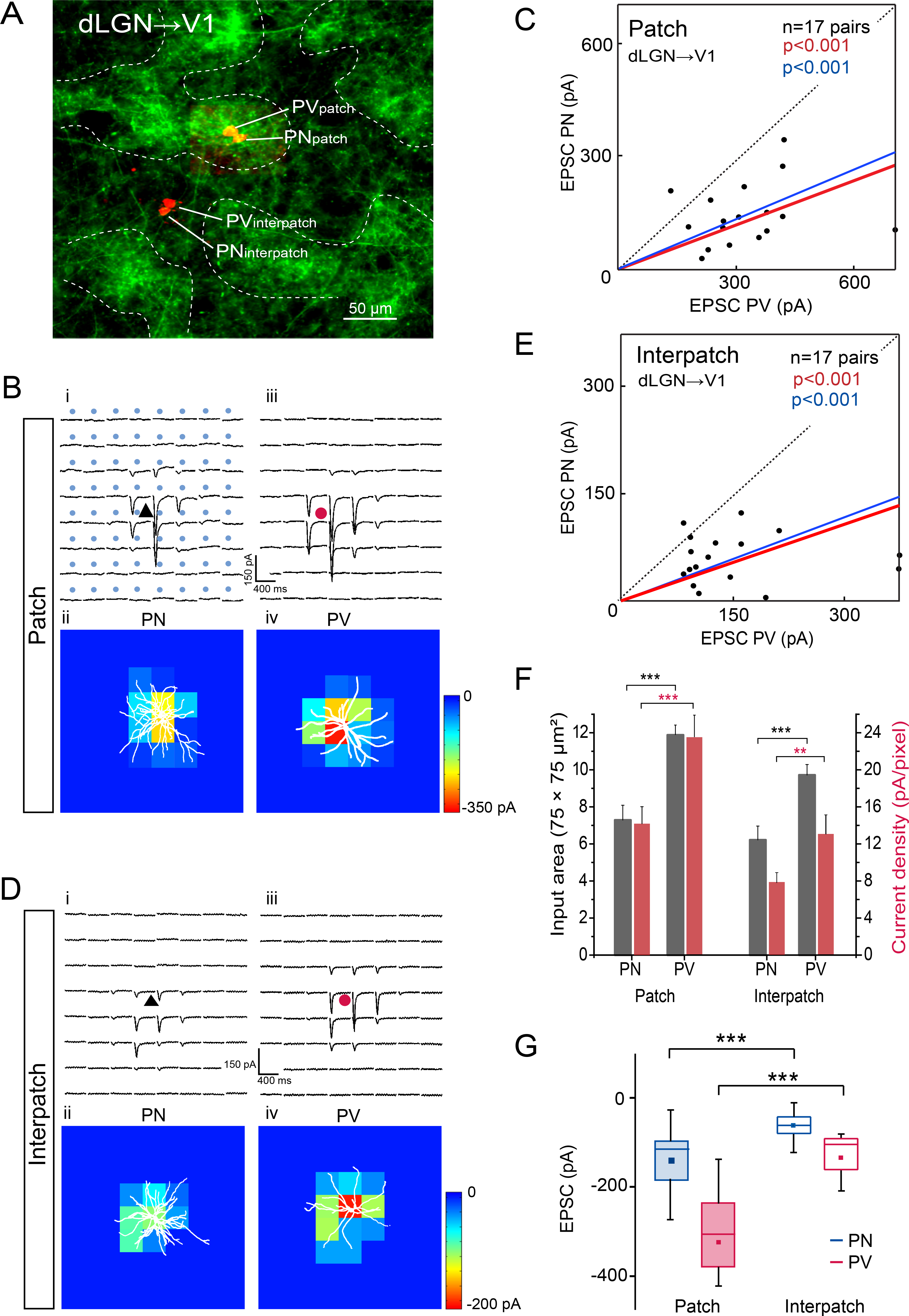
Tangential slices: sCRACM-mapping of dLGN→V1 input to L1 onto L2/3 PNs and PVs in patches and interpatches. (A) Z-stack showing ChR2-Venus labeled dLGN→V1 projections in L1 and Alexa 594 hydrazide-filled pairs of L2/3 PNs and PVs in patch and interpatch. (B-E) Recordings from PN (black triangle) and PV (red circle) in patches and interpatches obtained in the same slice. (Bi, Biii) Each trace represents average of EPSC_sCRACM_ (N=3) to laser stimulation (blue dots, 75×75 μm grid) of ChR2-expressing dLGN→V1 terminals. (Bii, Biv). Heatmaps of responses evoked at different locations of the dendritic arbor (Alexa 594 hydrazide-filled white profiles) of PN and PV shown in (Bi, Biii). (C) Each dot represents relative strength of dLGN input (summed pixels of significant EPSC_sCRACM_) of a pair of L2/3 PNs and PV in patch. Red line: mean slope from zero. Blue line: mean slope after normalizing currents to mean conductance. (A, Di-iv, E) Recordings from PNs and PVs in interpatch. Same conventions as in (B, C). (F) Distribution of dLGN input strength across dendritic arbor. Grey bars (number of pixels with non-zero EPSCs), PNs and PVs in patches and interpatches. Red bars, EPSC density (pA/μm^2^ per 75 μm x 75 μm pixel) in PVs and PNs of patches and interpatches. (G) Mean (dot) strength of dLGN→V1 EPSCs from PVs and PNs in patches and interpatches. (C, E, F, G) KS test. (\*\*\**p* < 0.001, \*\**p* < 0.01).

#### dLGN input in coronal slices

Although tangential slices are optimal for discriminating inputs to patches and interpatches, truncation of layers may favor stimulation of distal over basal dendrites. To control for potential preferences we obtained coronal slices of V1. To distinguish patches from interpatches we traced dLGN input to L1 with AAV2/9.CAG.ChR2.Venus. Photostimulation of ChR2-expressing dLGN inputs was performed in an 8 x 10 (mediolateral x dorsovental) 75 μm grid. Recordings were obtained from pairs of L2/3 PVs and PNs which were aligned with patches and interpatches. Although patchy dLGN Venus-labeled inputs were readily distinguishable at the slice surface, recordings from PNs and PVs from more than a single patch or interpatch in the same slice was challenging. Despite this caveat, which precluded recordings of PV/PN pairs from both patches and interpatches in the same slice, we found that in both module types EPSCs from PVs were lager (Figure 4A-C). Because ChR2 expression varied across animals and slices it was difficult to compare responses between experiments. But the range of response amplitudes suggests that the overall sizes of EPSCs in patches and interpatches is similar. This is consistent with the uniform distribution of dLGN inputs to L3/4 and suggests that the module-specific strength of activation we observed in tangential slices (Figure 3C, E, G) is due to the non-uniform distribution of L1 inputs.

**Figure 4.**
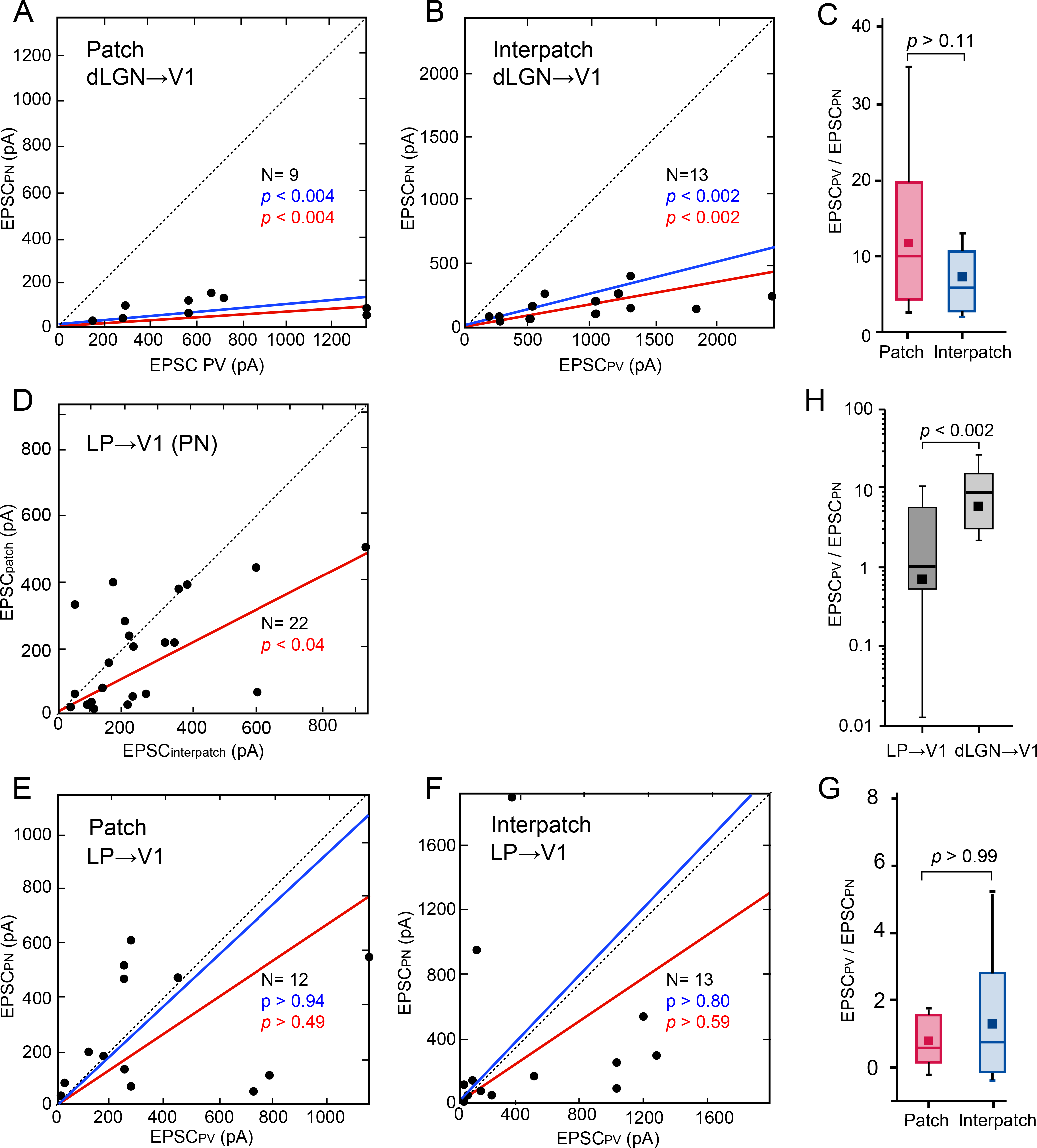
Coronal slices: sCRACM-mapping of dLGN→V1 and LP→V1 input to L2/3 PNs and PVs in patches and interpatches. (A, B) Relative strengths of EPSC_sCRACM_ from pairs (dots) of L2/3 PNs and PVs evoked by dLGN→V1 input. Recordings from patches (A) and interpatches (B) obtained in different slices. Mean slope of currents from zero (red), slope after normalization to conductance (blue). (C) Box plot of mean and median size of EPSC_sCRACM_ from PVs and PNs in patches and interpatches. (D) EPSC_sCRACM_ evoked by LP→V1 input to pairs of PNs in patches and interpatches recorded in the same slice. (E, F) EPSC_sCRACM_ evoked by LP→V1 inputs to pairs of PNs and PVs in patches and interpatches. Different pairs recorded in different slices. (G) Box plot EPSCPV/EPSCPN ratio in patches and interpatches. (H) Combined patch and interpatch box plots of EPSCPV/EPSCPN ratio in dLGN→V1 and LP→V1 pathways. (A-Wilcoxon signed-rank test.

#### LP input in coronal slices

Coronal slices were used for recording responses from L2/3 evoked by light stimulation of virally traced ChR2-expressing inputs from LP (Figure 1F). Patches were identified by viral tracing of dLGN input to L1 (Figure 1I). In recordings from the same slice we found that EPSCs from PNs were larger in interpatches than in patches (Figure 4D). This differs from the lack of modularity observed after stimulation of dLGN inputs in coronal slices (Figure 4A-C) and is consistent with the preferred innervation by LP of interpatches in L1 (Figure 1E-M). Light flashes in the LP-recipient L5A (Zhou et al., 2018) were ineffective in driving EPSCs from L2/3 PNs (data not shown). Thus, similar to patch-preferring dLGN inputs to L1 observed in tangential slices (Figure 3G), interpatch-preferring responses evoked by LP inputs in coronal slices are mediated through synapses in L1. Next, we compared the relative strengths of L1 LP-synapses in activating PNs and PVs in patches and interpatches in different slices. Unlike the strong bias for PVs observed by inputs from the dLGN (Figure 1G), we found that responses in patches and interpatches to LP inputs were matched, with a median EPSCPN/EPSCPV ratio in both compartments not significantly different than unity (Figure 4E-H). These results suggest that FFI in L2/3 produced through synapses in L1 is weaker in the LP→V1 than the dLGN→V1 pathway.

#### Feedback input from LM

Similar to thalamocortical pathways, input from LM→V1 terminated in periodic clusters of L1 where they largely overlapped with projections from the dLGN (Figures 2A). Photostimulation of ChR2-expressing LM→V1 feedback axons and comparing EPSCs from PNs and PVs in patches and interpatches of the same tangential slice, showed that responses in patches were larger (*p*<0.001, t-test) than in interpatches (Figure 5A, B, D, G), which correlated with the higher input density to L1 patches (Figure 2B). Different from the dLGN→V1 input, feedback from LM to patches evoked similar (*p* = 0.38, Wilcoxon signed-rank test) size EPSCs from PVs (529.5 ± 305.9 pA, n = 23) and PNs (598.9 ± 433.2 pA, n = 23) (Figure 5C, G). The even strength of LM input to V1 PNs and PVs in patches was also observed in the equal spread of activation across the dendritic tree and the matching current densities (Figure 5F). The spatial extent and current density in LM-activated patch-was larger than in interpatch-PNs (*p* < 0.01, two-sample t-test), whereas both measurements were similar for PVs (Figure 5F), suggesting that in the LM→V1 pathway the balanced activation in patches is due to stronger input to PNs. In contrast, responses from PVs (244 ± 186.4 pA, n=23) in interpatches were 1.9-fold larger (*p* < 0.001, paired t-test) than from PNs (130.1 ± 145.9 pA, n=23) (Figures 5E, G), an imbalance that was similar to that of dLGN inputs. Thus, balanced strength of input to PVs and PNs is a signature feature of higher order thalamic pathways, but not of top-down feedback, which is module-specific.

**Figure 5.**
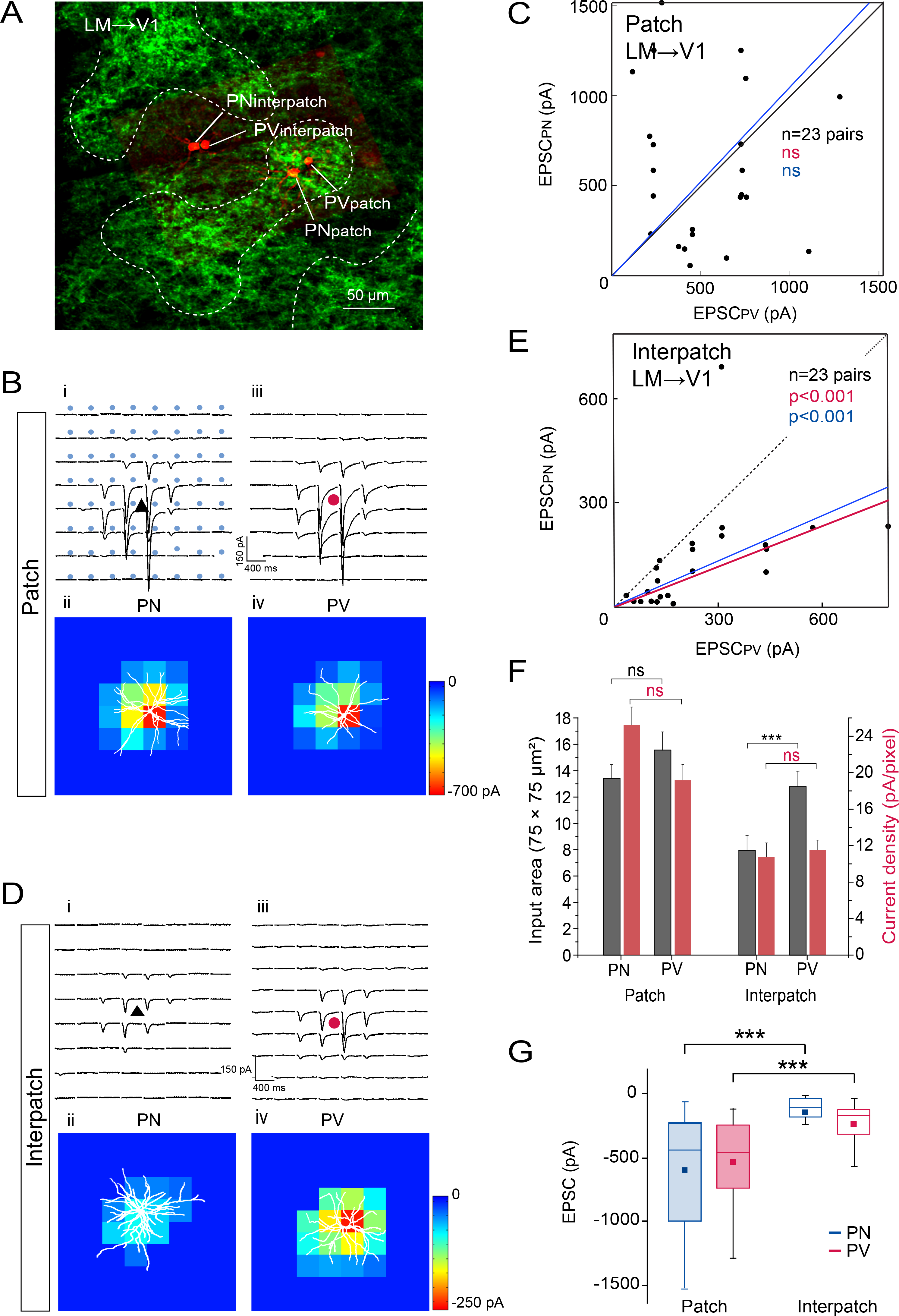
Tangential slices: sCRACM-mapping of LM→V1 input to L1 onto L2/3 PNs and PVs in patches and interpatches. Same conventions as in Figure 3. (A) Venus labeled LM→V1 projections in L1 and Alexa 594 hydrazide-filled pairs of L2/3 PNs and PVs in patch and interpatch. (B-E) Recordings from PNs and PVs in patches and interpatches obtained in the same slice. (Bi, Biii) Each trace represents average of EPSC_sCRACM_ to laser stimulation of ChR2-expressing LM→V1 terminals. (Bii, Biv). Heatmaps of responses from PN and PV shown in (Bi, Biii). (C) Dots represent relative strength of LM input to a pair of L2/3 PNs and PV in patch. (A, Di-iv, E) Recordings from PNs and PVs in interpatch. Same conventions as in (B, C). (F) Distribution of LM input strength across dendritic arbor. Grey bars (number of non-zero EPSC pxels), PNs and PVs in patches and interpatches. Red bars, EPSC/pixel density in PVs and PNs of patches and interpatches. (G) Box plots of strength of LM→V1 EPSCs from PVs and PNs in patches and interpatches. (C, E, F, G) Wilcoxon signed-rank test (\*\*\**p* < 0.001, ns=not significant).

### Long-range inputs to patches and interpatches drive distinct inhibitory subnetworks

The spatial clustering in the cell-specific strength of activation by inputs to L1 raised the question whether inhibition is non-uniformly mapped across the sheet of V1.

#### Clustering of GABAergic neurons

It has been reported that PV- and SOM-expressing GABAergic neurons in L2/3 of mouse V1 are clustered in the tangential plane and are radially aligned in L5 with subcortically projecting PNs (Ebina et al., 2014. Maruoka et al., 2017). To test this notion directly we cut tangential sections through V1 of PVtdT mice and found a striking clustering of tdT labeled fibers in L1 with a center-to-center spacing of ~120 μm (Figure S6). Clustering of PVtdT fibers was also present in surrounding cortex, most notably in LM, AL and primary auditory cortex (AUDp). To determine whether M2 and PVs are spatially registered, we stained tangential sections through V1 of PVtdT mice with an antibody against M2. We found that processes of PVs in L1-2/3 were strikingly clustered (Figure 6B-K). In the outer half of L1 (L1A, Figure 3A) the clusters contained mostly thick, smooth dendrites which overlapped with M2+ patches (Figure 6B-D). In contrast, in L1B and the top of L2/3 the PVtdT clusters switched to overlap with M2- interpatches. In L1B PVtdT clusters contained beaded dendrites and axons to which cell bodies and pericellular baskets, characteristic for basket cells (BC), were added in L2 (Figure 6A, E-K). The complementary mapping schemes were confirmed by the shifting distributions in PVtdT fluorescence intensities from patches (*p* > 10^−15^, KS) to interpatches (*p* = 3.42 x 10^−4^, KS) across the depth of L1-2 (Figures 6L, M). In separate counts we found that the PV cell density in the top 160 μm of L2/3 was 37% higher in interpatches (204/mm^2^) than in patches (128/mm^2^). To further support the module-selective mapping of PV axons we filled 8 PVs with biocytin. We found that in both patches (N=3) and interpatches (N=5) axons were largely confined to a ~100 μm-wide cluster and projections to the surround were sparse (Figure 6N). Notably, this was the case also for the projections parallel to the long axis of an obliquely cut, oval interpatch.

**Figure 6.**
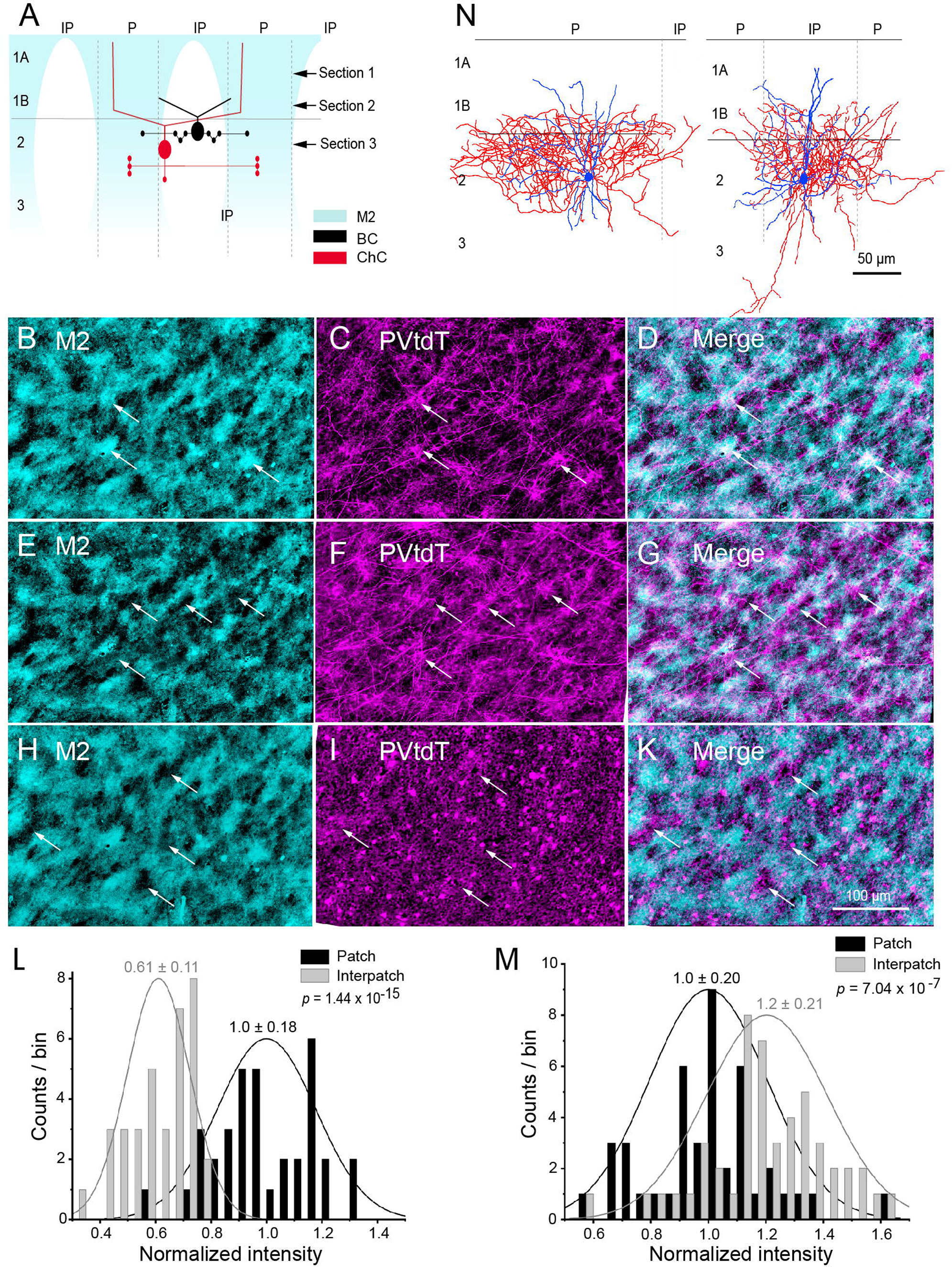
Spatial clustering of PV neurons in V1. (A) Diagram of coronal section through L1-3 showing M2+ patches (P), M2- interpatches (IP), and the preferred location of cell bodies, dendrites and boutons of PV-expressing basket (BC) and chandelier (ChC) cells. Arrows indicate positon of tangential sections (1-3) shown in (B-K). (B-D) Section 1 (A), immunolabeled M2+ patches (cyan) overlap (arrows) with PVtdT-expressing processes (magenta). (E-G) Section 2 (A), M2- interpatches (aligned to section in B) overlap (arrows) with PVtdT processes. (H-K) Section 3 (A), M2- interpatches (aligned to section in B) overlap (arrows) with PVtdT cell bodies, dendrites and boutons. (L) Normalized (to patches) fluorescence intensity at top of L1 (Section 1 showing stronger PVtdT expression in patches. (M) Normalized (to patches) fluorescence intensity in L2 (Section 3) showing stronger PVtdT expression in interpatches. KS test, mean ± SD (L, M). (N) Biocytin-filled L2/3 PV+ BCs (coronal plane) showing that dendrites (blue) branch in L1B-2. Axons (red) of cells in P (left panel) and IP (right panel) branch preferentially near the cell body with little spread into neighboring IP or P territories, respectively. The left panel shows a P cell with an asymmetrical axon arbor, which is mostly contained within P territory and shows sparse projections to IP. The elongated shape of the axonal arbor is caused by sectioning the oval-shaped P parallel to the long axis. PVs are rare in P, but branch density is similar to that of PVs in IPs.

Next, we asked whether the clustering of GABAergic neurons is subtype-specific. Most (>97%) types of GABAergic neurons express the vesicular GABA transporter (VGAT) (Uematsu et al. 2008). We therefore used VGAT-EYFP mice to look for patchy VGAT expression in V1. We found that VGAT expression in L1-2 was non-uniform but less discrete than the clustered projections from the dLGN (Figure S7A-F). Nevertheless, compared to unity of the shuffled patch/interpatch ratio, VGAT fluorescence intensity was significantly (*p* < 0.035) higher in interpatches, consistent with the clustering of PV (Figure 6M).

The phase shift of PVtdT and M2 between outer and inner parts of L1 suggested that this might be due to the differential distributions of chandelier (ChC) and BC dendrites in L1A and 1B relative to patches and interpatches (Figure 6A), similar to those found in mouse prefrontal cortex (Miyamae et al., 2017). To approach this question we compared the density of PVtdT boutons in cartridges apposed to Ankyrin G immunolabeled axon initial segments (AIS) (Blasquez-Lorca et al., 2015) in 75 μm-wide L2/3 ‘columns’ aligned with M2+ patches and M2- interpatches of PVtdT mice (Figure S8A-D). For analysis we selected radially oriented, tapered AISs of putative PNs. Consistent with the higher density of interneurons (Figure 6L) we found that the Ankyrin G-positive AIS density in interpatches was 21% lower (*p*<0.01, paired t-test) than in patches (Figure S8E), suggesting that the overall cell density across the cortical sheet is constant. Most notably we found that the length density of PV boutons in cartridges was 52% higher (*p*<0.01, paired t-test) in patches, which supports the notion that ChCs connections are preferentially aligned with patches (Figure 6A; S7E).

#### Subnetwork-specific FFI

Because FFI onto PNs depends on the strengths of excitatory long-range input to PVs and their inhibitory output (Isaacson and Scanziani, 2011), we performed whole cell recordings of unitary uEPSCs and uIPSCs from synaptically connected pairs of L2/3 PVs and PNs in V1 of PVtdT mice. To distinguish patches from interpatches we traced dLGN input to L1 with AAV2/9.CAG.ChR2.Venus. We first identified PVs by the expression of tdT and their non-adapting fast spiking properties and then searched for a neighboring PNs based on the triangular morphology and the regular spiking pattern (Figure 7A-C). Recordings from PNs were performed at holding potentials of −70 mV with pipettes containing a high [Cl^−^]. This enhanced the inward-directed (Luo et al., 2013), monosynaptic uIPSCs (Dong et al., 2004) (latency 2.4 ± 0.3 ms [patches n=22], 2.3 ± 0.3 ms [interpatches, n=26]), elicited by spikes (recorded in current clamp) from presynaptic PVs (Figure 7D, E). We found that uIPSCs and charge transfer (averaged across 50-150 repetitions) from PNs in interpatches (209 ± 149 pA, 2.79 ± 1.64 pC, n=22 pairs) were 5-7-fold larger (*p* < 0.001, two-sample t-test) than in patches (30 ± 15 pA, 0.54 ± 0.29 pC, n=26 pairs) (Figure 7F, G). Bath-application of Picrotoxin (50 μM) completely abolished uIPSCs in patches and interpatches (Figure 7D-F), demonstrating that responses were mediated via GABAA-receptors. Although PVs were less numerous in patches than interpatches, the probability of PVs→PNs contacts in patches (72%, 26 PNs/36 PVs) and interpatches (88%, 22 PNs/25 PVs) were similar (Figure 7H). These results show that the FFI subnetworks in patches and interpatches are not identical.

**Figure 7.**
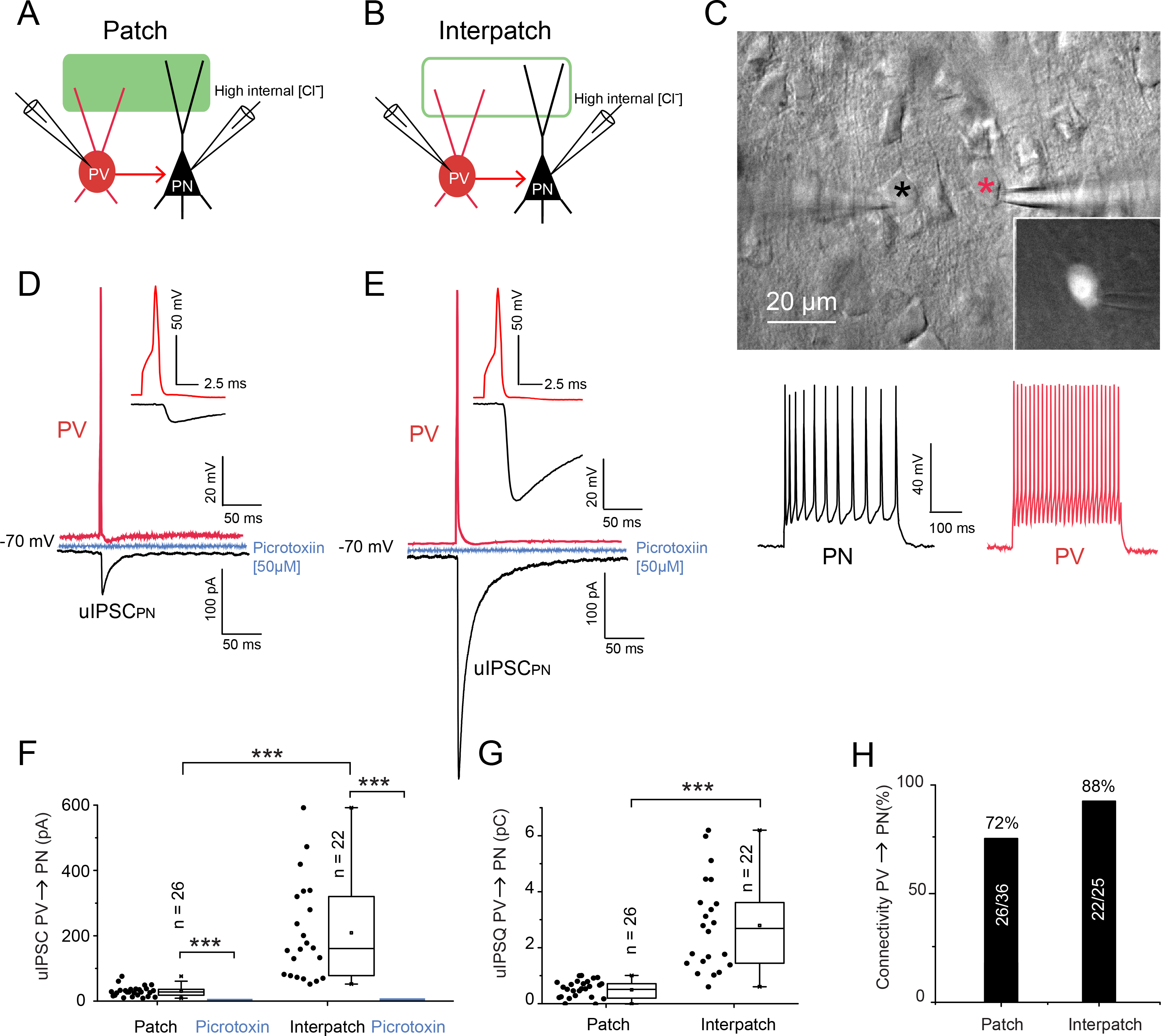
Distinct feedforward inhibition in patches and interpatches. A, B) Diagram of dual whole cell patch clamp recordings in coronal V1 slices of uIPSC in pairs of L2/3 presynaptic PV and postsynaptic PN aligned with ChR2.Venus-expressing dLGN→V1 patches (solid green) and interpatches (green outline) in L1. Recordings from PNs were made with high [Cl^−^] internal solution to shift equilibrium potential of IPSCs to 0 mV and record IPSCs at −70 mV as inward currents. Recordings from PVs were made with K-gluconate internal solution. (C) IR-DIC image showing paired recordings from L2/3 PN (black *, regular spiking shown below), and PV (red *, tdT expression shown in inset, fast spiking response below). (D, E) Recordings from synaptically connected PVs and PNs in patch (D) and interpatch (E). Spike fired by current injection in presynaptic PV (red trace). Voltage clamp recording of uIPSCs from postsynaptic PNs (black). Note that uIPSCs are larger in interpatches (E) than patches (D). uIPSCs are blocked by bath application of Picrotoxin (blue). The insets in (D) and (E) show that uIPSCs follow presynaptic spikes with delays consistent with monosynaptic connections. (F) Box plots of uIPSC recorded from PNs in patches and interpatches. Bath application of Picrotoxin abolished uIPSCs (\*\*\**p* < 0.001, two-sample t-test). uIPSCs in interpatches are larger (\*\*\**p* < 0.001). (G) The total unitary inhibitory charge (IPSQ) is larger (\*\*\**p* < 0.001, two-sample t-test) in interpatches than patches. (H) The connection probability between PVs and PNs in patches and interpatches is similar.

#### Subnetwork-specific E/I balance

Subnetwork-specific FFI differentially affect the mono- and polysynaptic excitation of PNs, which project back onto PVs and in turn may provide distinct feedback inhibition to PNs in patches and interpatches (Isaacson and Scanziani, 2011). In experiments separate from those shown in Figure 7, we measured the strength of uEPSCs and uIPSCs from reciprocally connected PN↔PV-pairs in patches and interpatches. The experimental procedures were similar to those in Figure 7, except that we also measured uEPSCs from PVs elicited by spikes in presynaptic PNs (Figure 8A-D). We found that uEPSC amplitudes and charge transfers from PVs in patches (42.2 ± 9.2 pA) and interpatches (29.1 ± 4.7 pA) were similar (*p* = ns, two-sample t-test) (Figure 8C-F, G, I). In sharp contrast, mean uIPSCs and uIPSQs recorded from PNs in interpatches (238.5 ± 45.9 pA, n =12 pairs) were 8.8-fold larger (*p* < 0.001, two-sample t-test) than in patches (27 ± 4.5 pA (n=12 pairs) (Figure 8C-F, H, J). Bath application of Picrotoxin (50 μM) completely blocked uIPSCs in patches (10/10) and interpatches (11/11) (Figures 8C, D). In the returning excitatory connections, bath application of DNQX (20 μM) completely blocked uEPSCs in patches (9/9) and interpatches (10/10), indicating that uEPSCs were mediated via AMPA receptors (Figures 8C, D). To estimate the relative strength of E and I we determined uIPSCs/uEPSCs ratios in patches and interpatches. In patches I/E was 0.89 ± 0.21 (n = 12 pairs) and differed significantly (*p* < 0.001, two-sample t-test) from 10.9 ± 2.5 (n = 12 pairs) in interpatches (Figure 8H). Similar I/E ratios were obtained for synaptic charge transfer (Figure 8J).

**Figure 8.**
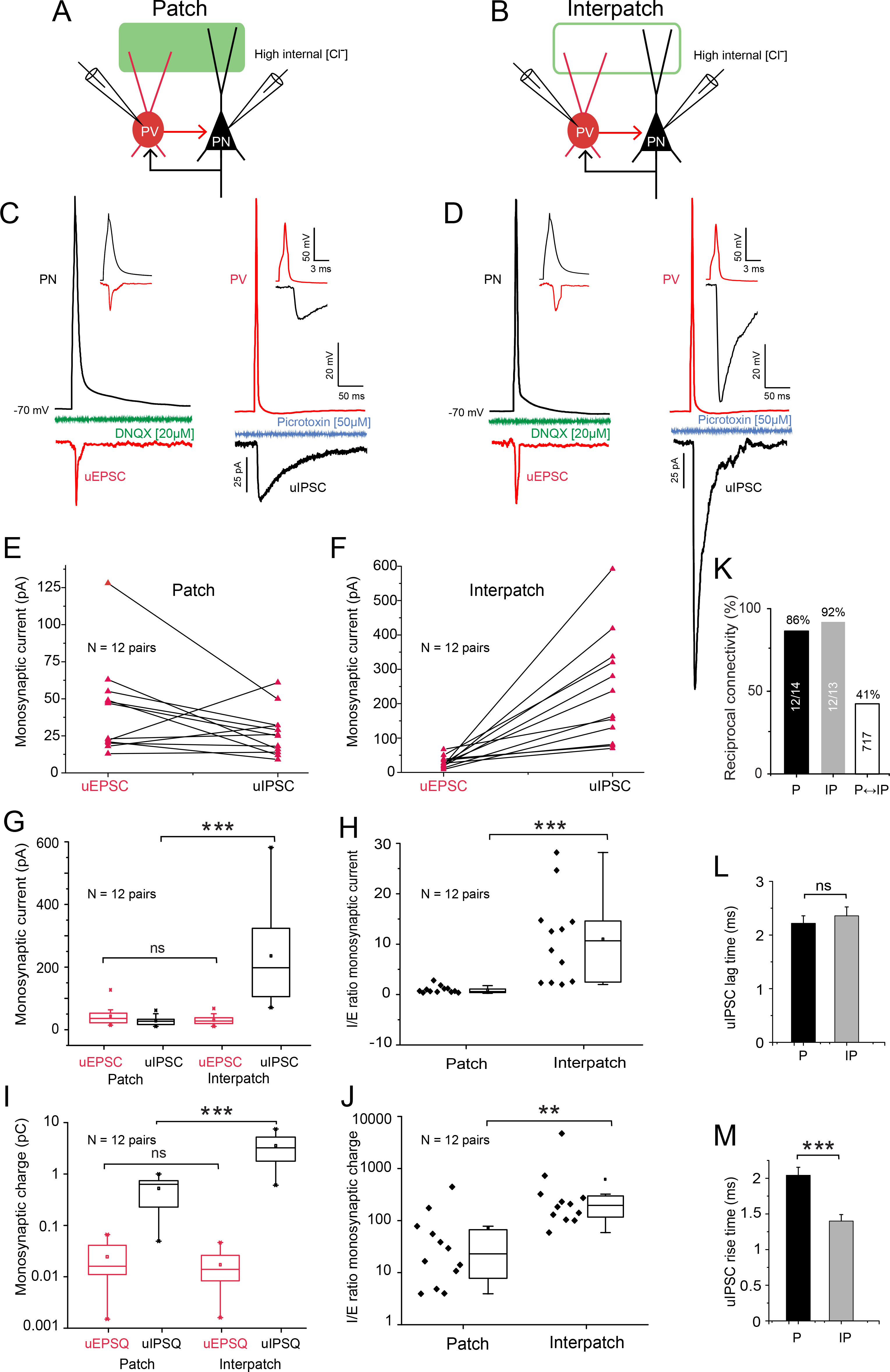
Stronger inhibition in interpatches than patches. (A, B) Diagram of recordings in coronal slices of reciprocal uIPSCs and uEPSCs in synaptically connected pairs of L/2/3 PNs and PVs aligned with ChR2.Venus-expressing dLGN→V1 patches (solid green) and interpatches (green outline) in L1. Recordings from PNs were made with high [Cl^−^] internal solution. Recordings from PV were done with K-gluconate solution in the pipette. (C, D) Monosynaptic uEPSCs (red trace) recorded from PV in response to a single spike (black trace) fired by L2/3 PNs in patch (C) or interpatch (D). Note that uEPSC in patches and interpatches have similar amplitudes and are blocked by DNQX (green C, D). In the reverse connection uIPSCs in interpatches (D) were larger than in patches (C), and were blocked by Picrotoxin (blue C, D). The insets in (C) and (D) show that in both directions PV→PN and PV←PN postsynaptic responses followed spikes with lag times indicating monosynaptic connections. (E, F) uEPSCs and uIPSCs recorded in pairs of PNs and PV in patches and interpatches. In comparing (E) and (F), note that in most pairs uIPSCs in interpatches are larger than in patches. (G) Average monosynaptic uEPSCs from reciprocally connected PV↔PN pairs are similar (ns, one-way ANOVA), whereas uIPSCs in interpatches are larger (\*\*\**p* < 0.001, one-way ANOVA). (H) I/E ratio of reciprocally connected PV↔PN pairs across patches and interpatches. Note that I/E balance in interpatches is tilted toward inhibition (\*\*\**p* < 0.001, two-sample t-test). (I) Average charge of uEPSCs and uIPSCs in patches and interpatches, showing that excitatory charge transfer at PN→PV contacts in patches and interpatches is similar, whereas the inhibitory charge transfer at PV→PN contacts in interpatches is larger (*** *p*<0.001, one-way ANOVA, Bonferroni correction for multiple comparisons). (J) I/E ratio showing that in reciprocally connected PV↔PN pairs in interpatches, uEPSCs are more strongly (** *p*<0.01, two-sample t-test) opposed by uIPSCs. (K) The reciprocal connectivity within patches and interpatches is 2.5-fold higher than between P and IP. (L) The onset latency of uIPSC in patches and interpatches is similar. (M) The rise time of uIPSCs in interpatches is faster than in P (\*\*\**p*<0.001, two-sample t-test).

In both patches and interpatches uIPSCs lagged uEPSCs by 2.21 ± 0.14 ms and 2.35 ± 0.16 ms, respectively (Figure 8L). Importantly, uIPSCs in interpatches showed significantly faster rise times (*p* < 0.001, two-sample t-test) than in patches (Figure 8M). The results show that although synaptic activation of PVs tracked that of PNs, the opposing inhibition of PNs is markedly stronger and faster in interpatches than in patches.

Because of the low density of PVs in patches, reciprocally connected PV↔PNs pairs (> 50 μm apart) were less common than in interpatches. Despite this difference the probability of mutually connected pairs in patches (81%) and interpatches (86%) was high (Figure 8K) and similar to PV→PN pairs (Figure 7H). In contrast only 33% PV↔PNs connections within a > 50 μm radius crossed the patch/interpatch border (Figure 8K). These findings are consistent with results showing that PVs are integrated into different subnetworks of PNs with similar response properties (Znamenskiy et al., 2018).

## DISCUSSION

We have found two interdigitating maps of M2+ patches and M2- interpatches in L1 of mouse V1 and show that PV-mediated inhibition of neighboring L2/3 PNs is significantly stronger in interpatches than in patches. We further show that each network is driven by distinct long-range inputs to dendrites in L1. While patches are the preferred targets of dLGN, LM and AL, interpatches receive inputs from the LP thalamus and PM. Although these inputs are module-specific the strength of synaptic activation of PVs and PNs in patches and interpatches is pathway-specific. Specifically, in patches dLGN input excites PVs more strongly than PNs, whereas both cells types are activated equally by feedback from LM. Similarly, in interpatches LM inputs to PVs and PNs are matched whereas LM input excites PVs more strongly. Together the results show that long-range inputs play a role in the E/I balance but suggest that the spike output of PNs is filtered by the activation threshold of PVs and their diverse strengths of inhibiting PNs in patches and interpatches. Although the efficacy of E/I coupling is thought to provide for module-specific tuning (Isaacson and Scanziani, 2011), the net impact on the output from patches and interpatches is shortening the integration window and exacting the selection of synchronous inputs (Gabernet et al., 2005). However, the opposing overall inhibition is strongest in interpatches, reducing response gain, increasing stimulus sensitivity (Atallah et al., 2012; Katzner et al., 2011) and improving the robustness of temporal frequency tuning in the interpatch module (Ji et al., 2015; Zhu et al., 2015).

### Patchy networks in L1 of V1

Finding interdigitating maps of 60-80 μm-wide clusters of thalamic and intracortical inputs in L1 was unexpected, given the salt-and-pepper organization of mouse V1 (Ohki and Reid, 2007). However, interdigitating microcolumns of subcortically and intracortically projecting L5 neurons, including clusters of L2/3 neurons and bundled apical dendrites with similar orientation and SF tuning, were recently found in mouse V1 (Kondo et al., 2016; Maruoka et al., 2017R; Ringach et al., 2016). It was puzzling, though, why L5 neurons were mapped hexagonally with a periodicity of 30-45 μm and showed an organization that differed from the quasi-rectangular lattice and the 120 μm periodicity of M2 patches we have found in L1 (Ji et al., 2015). In speculating about the alignment of infra- and supragranular maps we noted in tangential sections that upper layers occupy a ~25% larger area than infragranular layers (A. Burkhalter, unpublished results). Thus, to maintain topographic alignment across layers, ascending dendrites from L5 may be bundled proximally (Innocenti and Vercelli, 2010), form wider tufts distally and become targets of thalamocortical and intracortical connections in L1-2.

Clustered long-range projections to L1 are known from horizontal and feedback networks in primate, cat and mouse V1 (Ji et al., 2015; Martin and Whitteridge, 1984; Stettler et al., 2002). Here, we show that L1 projections from dLGN and the LP are also clustered in interdigitating maps. Different from the canonical core dLGN→V1 pathway to L3-4 (Bickford et al., 2015), input to L1 originates in the dLGN shell. The shell receives connections from direction-selective retinal ganglion cells and the superior colliculus and without prior cortical processing delivers orientation and direction selective signals (Bickford et al., 2015; Cruz-Martin et al. 2014; Roth et al. 2016) to patches in L1. Recordings from thalamocortical terminals have shown that the dLGN shell adds locomotion and saccade signals to visual responses and informs dendrites in L1 whether the speed of self-motion is matched to the visual flow of the environment (Roth et al., 2016). LP input to L1 derives from multiple subnuclei (Bennett et al., 2019), including the anterior portion whose projections we have traced most successfully. Similar to the dLGN shell, LP receives direction selective input from the retina and the superior colliculus (Allen et al., 2016; Zhou et al., 2017) but unlike dLGN shell projections, LP inputs to L1 are tuned to the mismatch of self-motion and visual flow and provide information about moving objects in the environment (Roth et al., 2016).

We found that intracortical feedback connections are clustered as well, but patch- and interpatch-projections are not cleanly sorted by their sources in the ventral or dorsal streams (Wang et al., 2012). Instead patch-projections originate from ventral-(i.e. LM) and dorsal-stream areas (i.e. AL), whereas another dorsal area, PM, preferentially targets the boundary region shared by patches and interpatches. This diversity suggests that the dorsal network branches into AL-dominated and PM-dominated sub-streams, perhaps similar to those in the primate occipito-parietal network specialized for visually guided actions and spatial navigation (Kravitz et al., 2011). The convergence of feedback from LM and AL suggests that patches multiplex inputs (Kampa et al., 2011) from functionally non-matching presynaptic neurons (Glickfeld et al., 2013; Marshel et al., 2012). Alternatively, responses of feedforward and feedback terminals may differ, suggesting that feedforward signals are transformed in higher areas, then returned for subtraction from bottom-up inputs to V1, which sends the error message back up the hierarchy to optimize sensory predictions (Zmarz and Keller, 2016).

### Subnetwork-selective targeting of PNs and PVs

The overlap of clustered thalamocortical and cortico-cortical projections to patches and interpatches suggests that the inputs target L1 dendrites of PNs and PVs whose cell bodies map to distinct X/Y coordinates in the layers below (Cruikshank et al., 2012; Johnson and Burkhalter, 1996; Miyamae et al., 2017). Recordings of EPSCs from L2/3 PNs and PVs elicited by dLGN and LM inputs support this interpretation, and in agreement with spatial clustering of connections, show that patch-inputs are stronger than interpatch-inputs. LP inputs to PNs show a similar structure-function relationship. Similar to the activation of cortical feedforward pathways (D’Souza e al., 2016; Yang et al., 2013) dLGN inputs to PVs were stronger than to PNs. This differed from the balanced LP and LM inputs to PVs and PNs, which is typical for intracortical feedback connections (D’Souza et al., 2016).

Synaptic inputs to PN and PV dendrites in L1 readily elicit spikes from cell bodies in the layers below (Cauller and Connors, 1994; Hu et al., 2010; Larkum et al., 2009). Our results show that the absolute and relative strengths of long-range inputs to PNs and PVs vary by pathway, patches (PV/PN ≈ 2:1) and interpatches (PV/PN ≈ 1:1), and demonstrate that the balance by which the network-stabilizing FFI tracks excitation (Xue et al, 2014) is pathway- and module-specific. Strong FFI in the dLGN→V1 pathway may select for synchronous inputs and enhance stimulus detection, whereas weaker FFI in the LP→V1 and LM→V1 pathways may broaden the integration window of convergent inputs and enhance the discriminability of stimulus features (Gabernet, 2005; Wang et al., 2010). Although FFI in patches and interpatches is proportional to the long-range input and the tuning properties of PNs and PVs (Hofer et al., 2011) the inhibitory effect of this input on PN spike output is stronger in patches than interpatches.

Consistent with the nonrandom connectivity in L2/3, but counter to areal uniformity (Kim et al., 2017), we have found that PV somata and terminals in L1-2 are spatially clustered (Ichinhoe et al. 2003; Maruoka et al., 2017; Znamenskiy et al., 2018). Similar to the spatial clustering of IPSC amplitude (Ebina et al., 2014) and consistent with clustering of visual response properties and strong inhibition between visually co-tuned PVs and PNs (Ji et al., 2015; Znamenskiy et al., 2018) the results show stronger local inhibition in interpatches than in patches. This may be due to larger, more proximal and/or the higher density of PV synapses onto PNs (Kubota et al., 2015; Stüber et al., 2015). Although the subcellular organization remains speculative, our results show that independent of the mode of activation by long-range inputs spikes from PVs in interpatches evoke at least 5-fold larger uIPSCs in interpatches than in patches. This strong locally generated inhibition may lower the gain of PN spike output and increase the sensitivity and robustness of responses to TF in interpatches (Atallah et al., 2012; Ji et al., 2015; Katzner et al., 2011; Zhu et al. 2015).

A possible role of the patch/interpatch organization is the selective tuning of visual responses of projection neurons. The most direct link comes from L5 PNs. Here, subcortically (i.e. pons, LP, superior colliculus, striatum) projecting L5B PNs overlap with PV clusters suggesting that they receive interpatch input onto thick dendrites in L1 (Kim et al., 2015; Maruoka et al. 2017). These cells are sensitive to high TF and low SF, properties we have found in L2/3 of interpatches (Ji et al. 2015; Kim et al., 2015). In contrast, L5A intracortically projecting (i.e. local V1, medial and lateral extrastriate visual areas) neurons which are excluded from PV clusters may receive patch input onto thin dendrites in L1 (Kim et al., 2015; Maruoka et al, 2017). The subset of Efr3a-Cre L5A neurons are sensitive to high SF which is the preferred property of L2/3 patch neurons (Ji et al., 2015; Kim et al., 2015). The relationship with patches and interpatches of the L2/3 PNs we have studied remains to be determined. Among the many types of L2/3 V1 neurons (Harris et al., 2018) few have dedicated projection targets and most are of the broadcasting type, projecting to multiple functionally distinct areas (Andermann et al., 2011; Han et al., 2018; Marshel et al., 2011) making a clean separation into patch and interpatch L2/3 neurons unlikely.

Besides the overlap of PV-positive BC dendrites with interpatches in the inner part of L1, different PV-dendrites reach the pial surface and preferentially branch in patches. The radial distribution of genetically labeled interneuron dendrites across L1 suggests that these processes belong to ChCs with cell bodies in L2/3 (Taniguchi et al, 2013; Tasic et al., 2018). The clustering of putative ChC dendrites is consistent with reports that AIS of cortico-cortically projecting L2/3 PNs are innervated in ~60 μm wide clusters (Blazquez-Llorca et al., 2015; Farinas and DeFelipe, 1991). Our findings confirm these results and show that the innervation density of L2/3 AIS in patches is higher than in interpatches, the opposite of what we have found in the projection density of PVs. L1 patches receive dense inputs from the dLGN, LM and AL, which when stimulated may recruit ChC-mediated FFI in layer 2/3 PNs (Woodruff et al., 2011). Unlike BC-mediated FFI, which regulates spike output from L2/3 PN by coordinating the dendritic integration of bottom-up and top-down inputs (Larkum et al. 2007; Larkum, 2013), ChC-mediated FFI by L1 input may suppress PN output at the AIS and cancel the error signal to higher visual areas for example during eye movements.

## STAR * METHODS

Detailed methods are provided in the online version of this paper and include the following:

- EXPERIMENTAL MODEL AND SUBJECT DETAILS

Animals
- METHOD DETAILS

Tracing of connections Immunostaining
Slice preparation
Subcellular Channelrhodopsin-2 assisted circuit mapping (sCRACM) Photostimulation
Recordings from synaptically connected pairs
- DATA ANALYSIS

Contour plots of patches and interpatches
Quantification of florescence intensity
EPSCs and IPSCs
Confocal imaging and neuron reconstruction
Statistics

## SUPPLEMENTAL INFORMATION

Supplemental information includes Supplemental Experimental Procedures and eight figures.

## AUTHOR CONTRIBUTIONS

P.B., R.D.D and A.B. designed the research. P.B. and R.D.D. performed the physiological and most of the anatomical experiments, with significant help from A.M., W.J. and A.B. All authors contributed to the data analysis. A.M implemented software. A.B., P.B. and R.D.D. wrote the paper with inputs from all authors.

## ACKNOWLEDGEMENTS

We thank Hongkui Zheng of the Allen Institute for Brain Science for the AAV2/1hSyn.tdTomato.WPRE.bGH, James Fitzpatrick and Dennis Oakley (Washington University Center for Cellular Imaging) and Katia Valkova for technical support. Thanks also to Tim Holy for the VGAT-ChR2-EYFP mice. Supported by National Eye Institute grants RO1 EY16184, RO1 EY022090, RO1 EY027383 and the McDonnell Center for Systems Neuroscience.

## DECLARTION OF INTERESTS

The authors declare no competing interests.

## SUPPLEMENTAL INFORMATION

**Figure S1. Intensity maps of dLGN→V1 and LP→V1 projections**. (A, B) Distribution of fluorescence intensity of M2tdT (A) and dLGN→V1 input (B) (same sections as in Figure 1A, B). Patches are outlined by white lines (top 33% quantile), interpatches by cyan lines (bottom 33% quantile). (C, D) Interdigitating maps of dLGN→V1 (C) and LP→V1 (D) projections to L1 (same section as in Figure 1I, K).

**Figure S2. Cholinergic axons preferentially innervate M2 patches**. (A, B) Tangential section through V1 showing double immunofluorescence of M2 patches (magenta) and ChAT axons (green) in L1. (C) Merge of (A) and (B). (D) Normalized (to patches) fluorescence intensity showing stronger ChAT expression in patches. KS test, mean ± SD.

**Figure S3. Development of M2 patches**. (A, B) Tangential section through V1 showing patchy expression of M2 in L1 of 4 day-old Chrm2tdT mouse (A). At higher magnification membrane-bound M2 expression in L2/3 shows rings of crossectioned dendritic bundles (arrows) occupying the spaces between unstained cells bodies (B). (C) Patchy pattern of M2 immunostaining in L1 of V1 of P10 C57BL/6J mouse.

**Figure S4. Recordings of L2/3 PVs and PNs in patches and interpatches**. (A-C) Confocal z-stack (200 μm) of tangential slice through L1-2 of V1 showing Alexa 594 hydrazide-filled pairs of PVs and PNs (red) aligned with Venus-expressing patch of LGN→V1 input to L1 (green) and Venus-negative interpatch. (B) Same slice as in (A) labeled with an antibody against M2. (C) Overlay of (A) and (B). (D) Alexa 594 hydrazide-filled interpatch-PVs and -PNs shown in (A) with apical dendrites branching preferentially in interpatches (unshaded regions). (E) Patch-PVs and -PNs shown in (A) with dendrites preferentially branching in patches (shaded regions). (F-J) Same format as (A-E), except that the Venus-expressing patches represent LM→V1 inputs. Note that similar to dLGN projections, LM inputs overlap with M2 patches (H).

**Figure S5. Time-lag of EPSC_sCRACM_ recorded from L2/3 PVs and PNs in tangential slices of V1**. (A, B) The time to peak after photo-stimulation of dLGN→V1 and LM→V1 inputs to L1 showing faster (\**p* < 0.05, two-sample t-test) responses in PVs (red) than PNs (grey).

**Figure S6. Clustering of PVtdT fibers in V1**. Tangential section though L1 (70 μm below pial surface) of V1 of PVtdT mouse, showing clusters of tdT labeled dendrites and axons. Arrows points to branched blood vessel. Lateromedial area (LM), anterolateral area (AL), rostrolateral area (RL), anteromedial area (AM), posteromedial area (PM), Mediomedial area (MM), restrosplenial agranular area (RSDagl), primary auditory area (AUDp), Somatosensory barrel cortex (SSp-bdf).

**Figure S7. Preferential VGAT expression in interpatches**. (A-D) Tangential sections through L1B-2 of V1 of VGAT-EYFP mouse. (A, B) Patchy dLGN→V1 inputs to L1 labeled by tracing with AAV2/1hSyn.tdTomato.WPRE.bGH (A) and corresponding contour map of patches (white lines) and interpatches (cyan lines) (B). (C-E) Patchy VGAT expression (C) and corresponding contour map (D) showing preferential expression in interpatches (cyan lines). (E) Merge of A and C shows interdigitating pattern. (F) Fluorescence intensity of VGAT-EYFP in interpatches relative to dLGN input to patches.

**Figure S8. Ankyrin G immunolabeled axon initial segments (AIS) of L2/3 PNs aligned with M2 patches are densely innervated by PVs**. (A) M2 immunolabeling (white) in parasagittal section through V1 of PVtdT (magenta) mouse. Patch (P). Interpatch (IP). (B) Same section as in (A) showing Ankyrin G immunolabeling of AIS (green) and M2 expression (white). Boxes indicate regions in L2/3 aligned with patches and interpatches in which PV innervation of AIS were counted. (C) Overlay of (A) and (B). (D) Confocal image showing putative contacts of PV boutons (magenta) with Ankyrin G labeled AIS (green). (E) Number of AIS in L2/3 underneath patches and interpatches. Length density of PV contacts onto L2/3 AIS in patches and interpatches. Paired t-test,** p < 0.01.

## Supplemental Experimental Procedures

### Animals

Experiments were performed on male and female, C57BL/6J, PV-Cre (Bg.129P2-Pvalb^tm(cre)Arbr^/J) × Ai9 (Gt[ROSA]26Sor*tm9(CAG-tdTomato)Hze*), Chrm2-tdT-D knock-in mice (Bg6.Cg-Chrm2^tm1.1Hze^/J), Chrm2 M2R-/- (B6N.129S4(Cg)-Chrm2^tmJwe^/J) and VGAT-ChR2-EYFP (B6.Cg-Tg(Slc32a1-COP4*H134R/EYFP)8Gfng/J) mice. For anatomy we used 4-10 day old and >46 day old animals. Slice recordings were done in 34-46 day-old animals. Thalamocortical and intracortical connections to V1 were visualized by axonal tracing with AAV. sCRACM mapping of long-range input to PNs and PVs and recordings of uEPSCs and uIPSCs between synaptically connected pairs of PNs and PVs were performed in acute slices of V1. All experimental procedures were approved by the Institutional Animal Care and Use Committee at Washington University.

### Tracing of connections

Connections were traced anterogradely by intracerebral injection of AAV2/9.CAG.ChR2.Venus.WPRE.SV40 (Vector Core, University of Pennsylvania), AAV2/1.hSyn.EGFP.WPRE.bGH (Vector Core, University of Pennsylvania) and/or AAV2/1hSyn.tdTomato.WPRE.bGH (Allen Institute) in 18-20 day-old or 8-12 week-old mice. Animals were anesthetized with Ketamine/xylazine (86 mg.kg^−1^/13 mg.kg^−1^, IP). Analgesia was performed with Buprenorphine (0.1 mg/kg, SQ). The eyes were protected with ophthalmic ointment. All surgical procedures were performed in a stereotaxic apparatus. Injections (46 nl) were made with glass pipettes (tip diameter 15-25 μm) connected to a Nanoject II injector. Stereotaxic injections were made into the (in mm): dLGN (2.35 posterior of bregma, 2.15 lateral of midline and 2.55 below the pial surface), LP (1.85 posterior of bregma, 1.25 lateral of midline and 2.65 below the pial surface), higher visual cortical lateromedial area, LM, (1.4 anterior to transverse sinus, 4.1 lateral to midline, 0.3- 0.5 below the pial surface) and posteromedial area, PM, (1.9 anterior to transverse sinus, 1.6 lateral to midline, 0.3-0.5 below the pial surface). Postsurgical survival was 2-3 weeks.

### Immunostaining

Mice were overdosed with Ketamine/xylazine (170mg.kg^−1^) and transcardially perfused with heparinized phosphate-buffered saline (PBS), followed by 1% paraformaldehyde (PFA). Brains were extracted from the skull, the left cortical hemisphere was removed, flat mounted, postfixed overnight in 4% PFA and cryoprotected in 30% sucrose. Tangential or parasagittal sections were cut at 40 μm with a freezing microtome. Sections were washed in 0.1 M phosphate buffer (PB), treated in blocking solution containing 10% normal goat serum (NGS), and 0.1% Triton X-100 in PB. Immunolabeling was performed by incubating sections for 48 hours at 4ºC with primary antibodies against M2 muscarinic acetylcholine receptor (1:500 rat monoclonal, MAB367 Millipore) or anti-Ankyrin G (1:1000, mouse monoclonal, clone N106/36, NeuroMab). After washing, sections were treated with Alexa Fluor 647-labeled goat anti-rat IgG secondary antibody (1:500 in 10% NGS; A21247 Invitrogen) or donkey-anti-mouse-Cy5 (1:500, 715-175-151, Jackson Immuno Research). Cholinergic fibers were identified with an antibody against choline acetyl transferase (1:1000, goat anti-ChAT, Millipore AB144P), detected with a biotinylated donkey-anti goat secondary antibody (1:200, Millipore AP180B) and visualized with NeutrAvidin Oregon Green 488 (1:400, ThermoFisher A6374). Sections were mounted onto glass slides, coverslipped in PB or Aqua Poly/Mount (Polysciences) and imaged under a fluorescence microscope (Nikon 80i) equipped with a CCD camera (CoolSnap EZ, Roper Scientific or Infinity3S-URM, Lumenera). Confocal imaging was performed with an Olympus Fluoview (FV1200) microscope. The specificity of the M2 primary antibody was validated in C57BL/6J-M2^-^/^-^mice (Gomeza et al., 1999) in which we saw no detectable staining.

### Slice preparation

Slices of V1 were prepared from 34-46 days-old virus-injected mice. The slices were either cut in the tangential or the coronal plane. Tangential slices were optimally suited for identifying repeating clusters of ChR2.Venus-labeled thalamocortical and intracortical inputs to L1 and preserving the complete dendritic arbors of PNs and PVs. Mice were decapitated under isoflurane (2% in oxygen) anesthesia. The brain was rapidly removed from the skull and submerged in ice-cold cutting solution aerated with 95% O2/5% CO_2_ containing (in mM): 240 sucrose; 2.5 KCl, 1.25 NaH_2_PO_4_, 2.1 NaHCO_3_, 7 MgCl_2_, 0.5 CaCl_2_, 10 glucose, adjusted with NaOH to pH 7.35. Next, the cerebellum and anterior third of the brain were removed. With the cut rostral surface towards the base, the lower part of the brain was resected parallel to the surface of V1 and the tissue block was mounted with the cut-side down onto the specimen plate. Single tangential slices (350 μm) were cut in ice-cold cutting solution on a Vibratome (Leica VT 1200). Coronal slices were prepared as described previously (Yang et al., 2013). Slices were kept in a holding chamber in which they were submerged in oxygenated artificial cerebrospinal fluid (ACSF; containing [in mM]:125 NaCl, 2.5 KCl, 1.25 NaH_2_PO4, 25 NaHCO_3_, 1 MgCl_2_, 2 CaCl_2_, 25 glucose, pH 7.35) for 1 hour at 32ºC before transferring them to the recording chamber maintained at room temperature (22-24ºC).

Clustered projections were also readily identified in coronal slices which are optimal for preserving connections across layers. The procedures of preparing coronal slices were identical to those for tangential slices, except that the injected hemisphere was mounted with the coronally cut frontal surface down and serial slicing was from the posterior pole.

### Subcellular Channelrhodopsin-2 assisted circuit mapping (sCRACM)

For recording, slices were transferred to a submersion chamber mounted on the stage of a modified upright fluorescence microscope (Nikon Eclipse FN1) equipped with a CCD camera (Retiga-2000C; Qimaging). Slices where perfused (1.5 ml/min) with recirculating oxygenated ACSF (22-24ºC). Whole cell patch clamp recordings were obtained from pairs of tdT-expressing PVs (identified with fluorescence optics) and nearby (within < 40 μm) unlabeled PNs cells (identified with DIC-IR optics) in L2/3 of V1. Cell pairs were either in-register (i.e. within patches) or out-of-register (i.e. within interpatches) with ChR2.Venus-labeled patches of axons projecting from the dLGN, LP or area LM and terminating in L1 of V1. For recording in tangential slices, neurons were approached from the cut surface of the slice, which was mounted with the pial surface facing down. In tangential and coronal slices, recordings were made 30-120 μm below the surface of the slice. Electrical signals were sampled at 10 kHz by Multiclamp 700B amplifiers (Molecular Devices), digitized (NI USB 6363; National instruments) and acquired using Matlab-based (MathWorks, Natick, MA) Ephus software (Suter et al., 2010). Electrodes were pulled from borosilicate glass capillaries (G150F-4, Warner Instruments). Pipettes were filled with (in mM): 128 potassium gluconate, 4 MgCl_2_, 10 HEPES, 1 EGTA, 4 Na_2_ATP, 0.4 Na_2_GTP, 10 sodium phosphocreatine, 3 sodium L-ascorbate, 0.02% Alexa 594 hydrazide (Invitrogen), pH 7.25, 290 mOsm. The pipette resistance was 3-5 MΩ. The liquid junction potential was not corrected. The seal resistance was > 2 GΩ. Recordings with access resistance of > 20 MΩ were excluded from the study. Neuron type was assessed by recording spiking patterns (i.e. fast for PVs, regular for PNs) in response to 300 ms pulses of hyperpolarizing and depolarizing current in current clamp mode. For sCRACM mapping (Petreanu et al., 2009), EPSCs were recorded in voltage clamp at a holding potential of −70 mV, with tetrodotoxin (1 μM) and 4-aminopyridine (4-AP; 100 μM; Tocris Bioscience) in the bath to block action potentials and fast repolarizing potassium currents, respectively. After recording, slices were fixed in 4% PFA, cleared in 10% sorbitol, mounted on glass slides in Aqua Poly/Mount, imaged under a confocal microscope (Olympus, Fluoview FV1200) and reconstructed in 3D, using ImageJ.

### Photostimulation

Photostimulation was performed with a blue laser (473 nm; CrystaLaser). The light was reflected by a fixed set of mirrors onto galvanometer scanners (Cambridge Scanning) that controlled beam position. The light then passed through an air objective (4 PlanApo, NA 0.2; Nikon), which at 0.25 mW/cm^2^ laser power formed a beam at half maximal intensity with a diameter of ~20 μm in the specimen plane. The durations and intensities of the light pulses were controlled with a Pockels cell (ConOptics) and a shutter (LS6, Uniblitz). Because the proportion and labeling intensity of ChR2-expressing axons varied across slices and animals, the laser power (0.25–1 mW/cm^2^) was adjusted in every slice to evoke EPSC. The laser power was constant for all recordings made that day. Recordings in V1 were performed from pairs of nearby PNs and PVs in patch and interpatch regions of L2/3. Each trial consisted of 100 ms baseline, followed by the photostimulus (1–2 ms) and 300 ms of response. Photostimulation was performed in an 8×8 grid in which individual points were spaced 75 μm apart and the grid was aligned with the recorded soma at the center. For mapping in coronal slices, one side of the grid was aligned with the pial surface. The stimulation sequence was pseudorandom allowing maximal intervals between nearby stimulation sites. sCRACM maps were generated from 3-5 repetitions per neuron.

### Recordings from synaptically connected pairs

To examine the presence and strength of synaptic connections between PVs and PNs in patches and interpatches we recorded from synaptically connected pairs in superficial L2/3 of V1. Patchy projections to L1 were identified as clusters of Venus-expressing axon terminals, labeled by tracing dLGN→V1 inputs with AAV2/9.CAG.ChR2.Venus. Recordings were obtained from cell pairs (< 40 μm apart) aligned with patches or interpatches in coronal slices. Responses in PVs were recorded with pipettes (4-6 MΩ resistance) filled with (in mM): 128 potassium gluconate, 4 MgCl_2_, 10 HEPES, 1 EGTA, 4 Na_2_ATP, 0.4 Na_2_GTP, 10 sodium phosphocreatine, 3 sodium L-ascorbate, pH 7.25, 290 mOsm. For recording PNs, pipettes were filled with a high [Cl^−^] solution containing (in mM): 145 KCl, 10 HEPES, 5 NaATP, 0.2 NaGTP and 5 EGTA, pH 7.3, adjusted with KOH, 285 mOsm. Under these conditions the reversal potential of IPSCs is 0 mV and at −70 mV holding potential the currents flow inward (Luo et al., 2013). Monosynaptic uIPSCs were recorded by holding PNs at −70 mV. uIPSCs were elicited by triggering single action potentials from presynaptic PVs with 2 ms depolarizing current pulses. The same stimulation/recording paradigm was used for eliciting and measuring uEPSCs, except that spikes were elicited from PNs and responses were recorded from PVs. Responses were averaged across 50-150 repetitions at 0.5 Hz. The error due to the liquid junction potential was not corrected. Access resistance was monitored throughout the experiment. Cells whose series resistance was >20 MΩ or varied >25% for the duration of the experiment were excluded from the analysis. Series resistance errors were not compensated. To block spontaneous polysynaptic NMDA receptor-mediated excitatory currents, CPP ((RS)-3-(2-carboxypiperazin-4-yl)-propyl-1-phosphonic acid, 50 μM, Tocris) was applied in the bath. uIPSCs were blocked by bath application of the GABA_A_-receptor antagonist Picrotoxin (50μM, Tocris). uEPSCs were blocked by bath application of the AMPA-receptors antagonist DNQX (6,7-dinitroquinoxaline-2,3-dione, 20 μM, Tocris). After recording, the slices were fixed in 4% PFA, mounted on glass slides, cleared in 10% sorbitol and Alexa 594 hydrazide-filled neurons were imaged under the confocal microscope.

### Data analysis

#### Contour plots of patches and interpatches

Automatic patch/interpatch definition followed the general procedures of Sincich and Horton (2005). Fluorescent images of spatially clustered M2 expression or virally traced projection patterns in L1 of V1 were high-pass filtered using an 80 μm filter radius. Images were then blurred using a circular averaging filter of 30 μm radius, with the ‘fspecial’ function in Matlab. All pixels in the resulting images were then divided into six intensity quantiles. The top two quantiles were considered to be patches and the bottom two interpatches. For statistical testing, images in matching fields of view were analyzed. Images were downsampled to have a pixel area of 150 μm^2^ each. A permutation test was then performed by shuffling fluorescent image pixels within the image and determining the ratio of resulting average patch intensity to average interpatch intensity, maintaining the original patch/interpatch borders derived from M2 or viral tracings. Patch/interpatch ratios in the original image outside the 95% bounds of the randomized distribution from 100,000 shuffling iterations were considered significant deviations from a 1:1 patch/interpatch intensity ratio.

#### Quantification of fluorescence intensity

The intensity of immunofluorescence of M2, fluorescently labeled projections tagged by viral axonal tracing, and fluorescence of transcribed tdT and EGFP genes were quantified in images of tangential sections through L1, acquired with a CCD camera (Lumenera Infinity3S-URM) and Metamorph NX2.0 software (Molecular Devices). Gray scale images were opened with Image J, background subtracted to correct for global non-uniformities in brightness and overlaid with contour maps of fluorescence intensity determined by a custom Matlab script. Pixel values in patches and interpatches were measured at multiple sites, normalized to the mean brightness of patches, binned and plotted as counts of normalized fluorescence intensity. Statistical comparisons of intensity distributions were made using the KS test.

#### EPSCs and IPSCs

The amplitude of significant responses was >4 times the SD of the baseline. Individual pixel values of sCRACM maps were computed from the mean EPSC amplitude in a 75 ms response window after the photostimulus. For each neuron, maps were averaged across 3-5 repetitions. These averages represent synaptic charge transfer. Because the responses were dominated by the current amplitude and small long-lasting currents were negligible, we have adopted the simplification introduced by Petreanu et al., (2009) and represent responses in pA instead of Coulomb. The charge value for each pixel in a 75 ms window was calculated using custom Matlab software. EPSC amplitudes were measured with reference to the soma at the center. To display the scaled magnitudes and spatial distributions of thalamocortical and intracortical inputs from LM to PNs and PVs, maps for each cell class were peak normalized within individual slices and displayed as heatmaps. Comparisons of inputs to PNs and PVs were made by plotting the average responses from pairs of PNs and PVs within layer 2/3 of the same slice and in the same patch or interpatch module. Thalamocortical and intracortical inputs to PNs and PVs recorded in the same layer and same slice were plotted against each other and the relative strengths of excitation was assessed by plotting the mean slope from zero. uIPSC lag time was calculated as the time delay from the onset of the presynaptic PV spike to the onset of the uIPSC recorded from the postsynaptic PNs. uIPSC rise time was measured as the delay between response onset and the peak.

#### Confocal imaging and Neuron reconstruction

Alexa Fluor 594 hydrazide-filled neurons were reconstructed *posthoc* and their location in Venus-expressing patches of dLGN or LM inputs determined by imaging under a confocal microscope (Olympus, Fluoview FV 1200), using a 30x silicone oil (UPLSAPO, 1.05 NA, Olympus) objective. Twelve bit 1024 x 1024 pixel images were taken at 1.5x digital zoom. Z-stacks were acquired at 0.80 μM/section (Nyquist volume: 1.6 μM) across the thickness of the slice. Multicolor scanning was done in sequential and frame-by-frame mode. The images were acquired in separate high sensitivity detector channels for each fluorophore. The signals were acquired and averaged by Kalman’s method to increase signal/noise ratio. The neurons were then traced and reconstructed by using the ‘Simple Neurite Tracer’ Plugin of Fiji (ImageJ). PNs were identified by the presence of dendritic spines, whereas PVs have aspinous, beaded dendrites.

Recorded neurons were filled with biocytin (3 mg/ml) which after fixing slices in 4% PFA was visualized by an ABC reaction and intensification of the reaction product with AgNO3 and HAuCl2 (Yang et al., 2013). Filled neurons were reconstructed under a 40x oil objective using Neurolucida (MicroBrightField).

The innervation density of PN-AISs by PVtdT expressing boutons was determined by confocal imaging with a 100x oil immersion objective of z-stacks (0.2 μm step size) in ROIs (65×135μm) aligned with M2+ patches and M2- interpatches. To minimize contamination by Ankyrin G-expressing AIS of interneurons we focused the analysis on tapered, vertically (±3° relative to the pial surface) descending profiles. Appositions between boutons and AIS were scored as contacts if there was no detectable gap between pre-and postsynaptic elements and their association remained stable under image rotation.

#### Statistics

Statistical analyses were performed using Origin 9.1 (Origin Laboratory) or customized Matlab software. Normality was assessed using the Shapiro-Wilk normality test to select between parametric and nonparametric tests. Comparisons between two groups were performed with two-tailed Student’s *t*-test. Neighboring neurons that were recorded sequentially were considered pairs and subjected to a paired *t*-test. For comparisons across more than two groups, data were analyzed using one-way ANOVA followed by Bonferroni’s *post hoc* analysis to correct for multiple comparisons. For data with non-normal distribution, nonparametric KS test was used. Significance was *p* < 0.05. Data are mean ± SEM, except when otherwise indicated as mean ± SD. Box plots mean ± SD.

