## Supplementary figures and images for "Spatial clustering of inhibition in mouse primary visual cortex"

### Fig S1

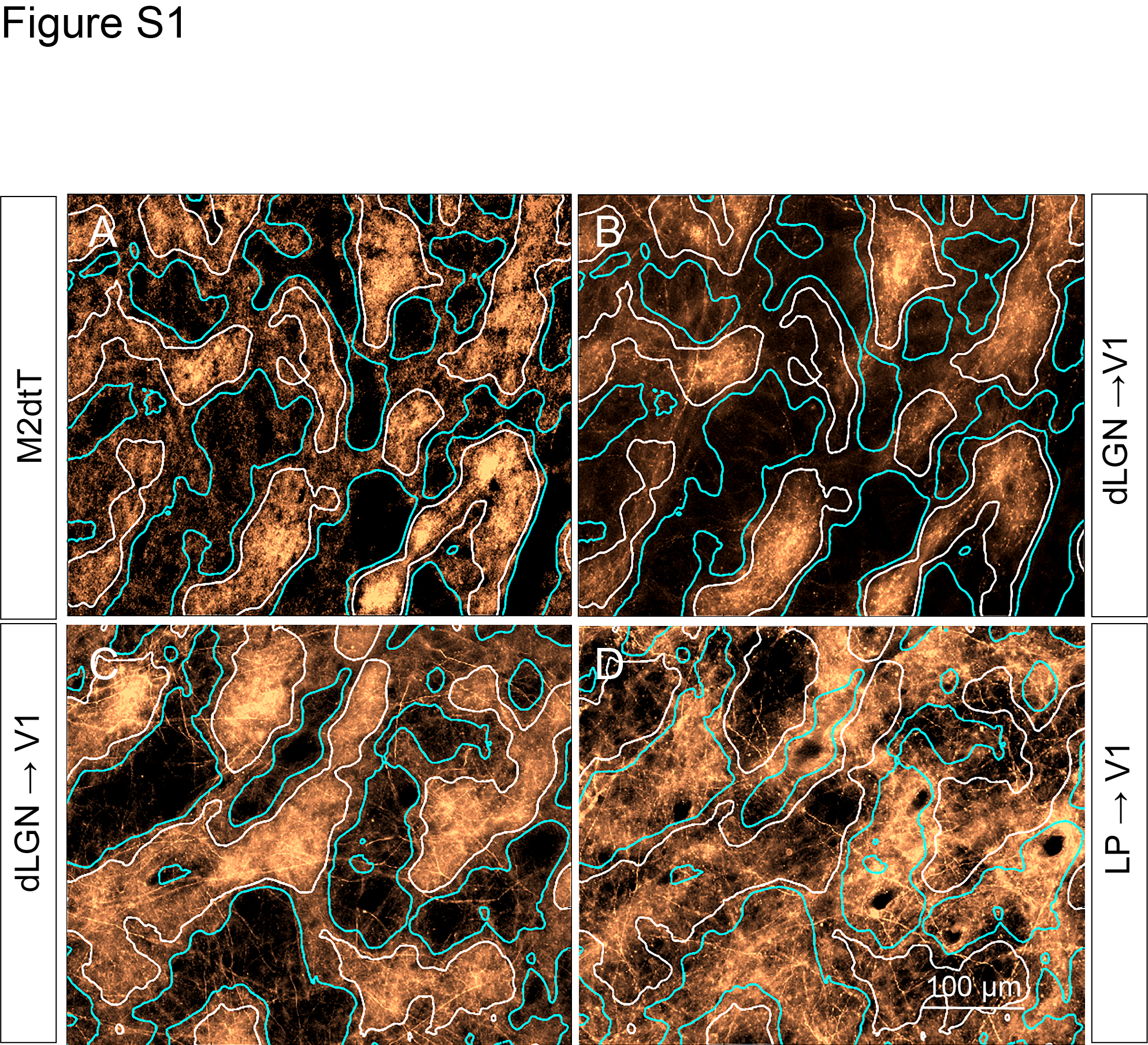

### Fig S2

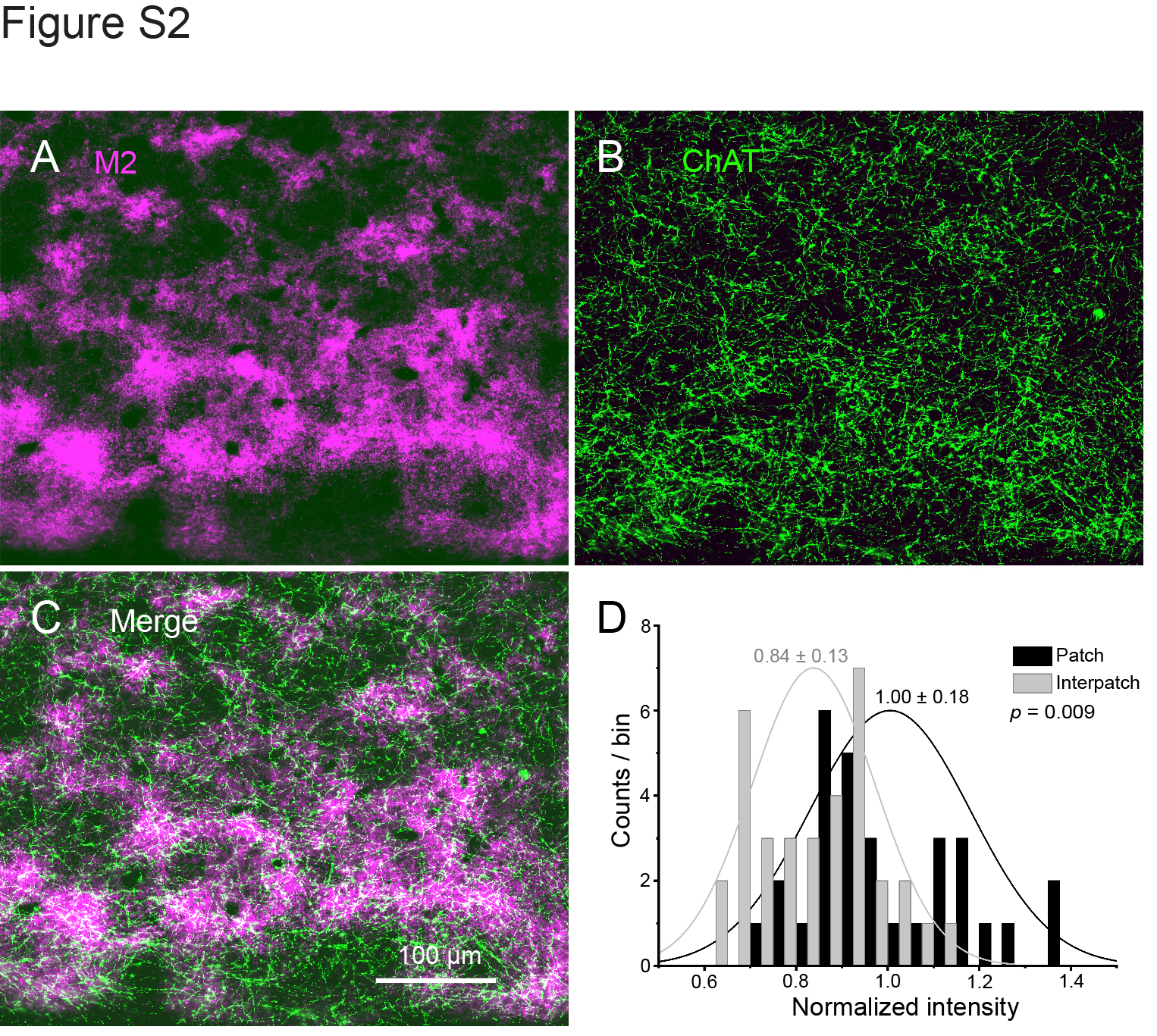

### Fig S3

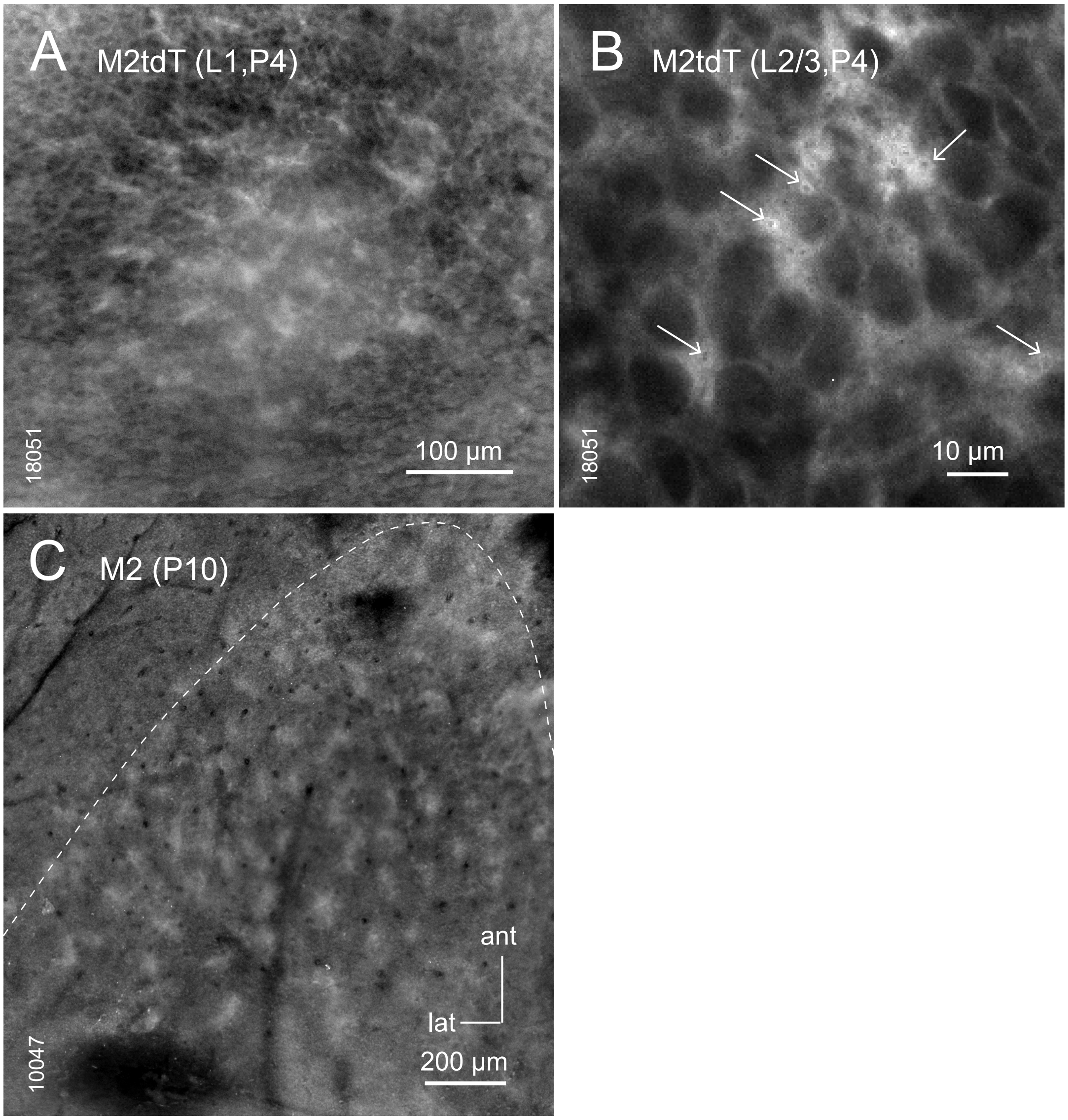

### Fig S4

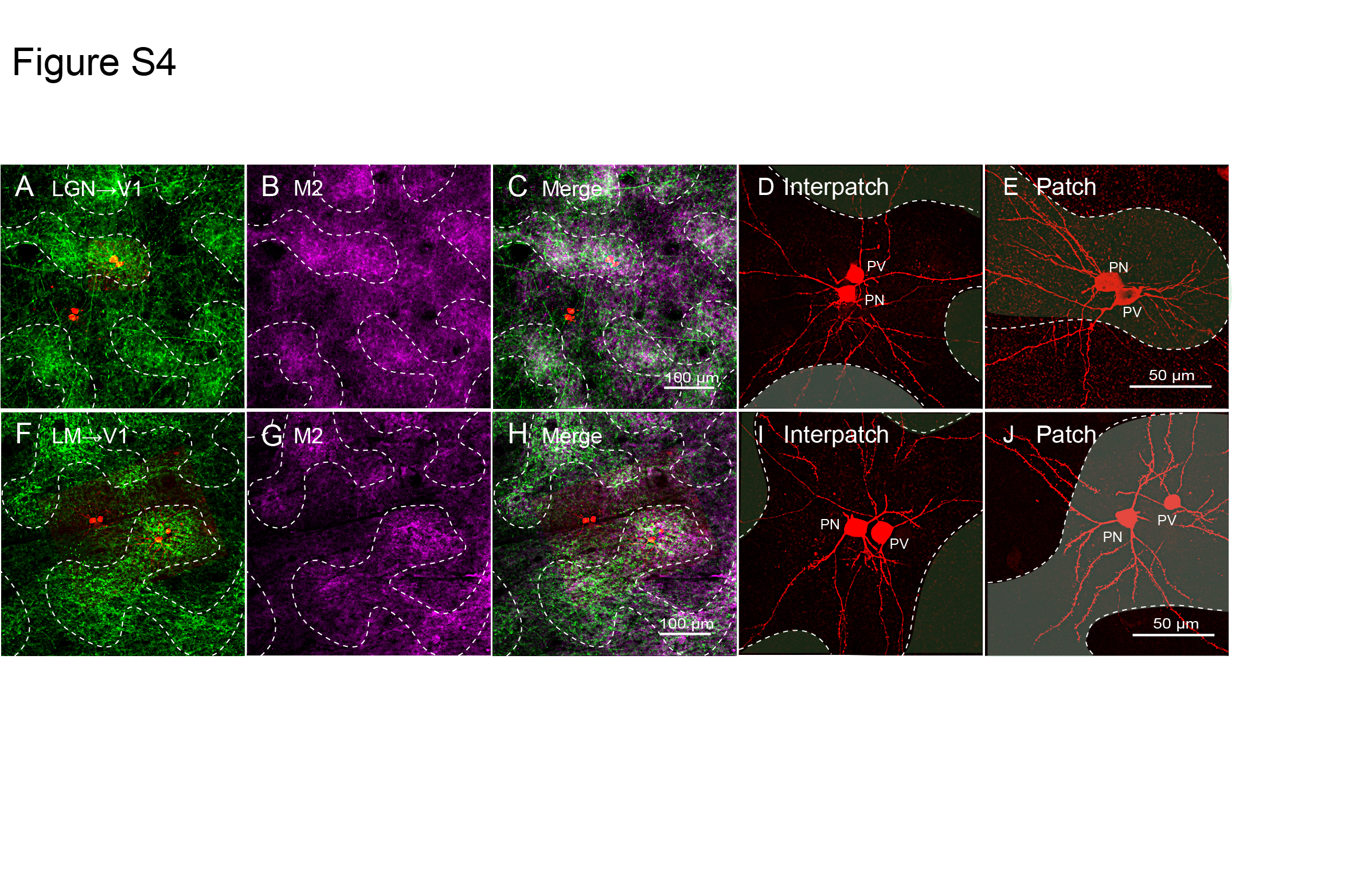

### Fig S5

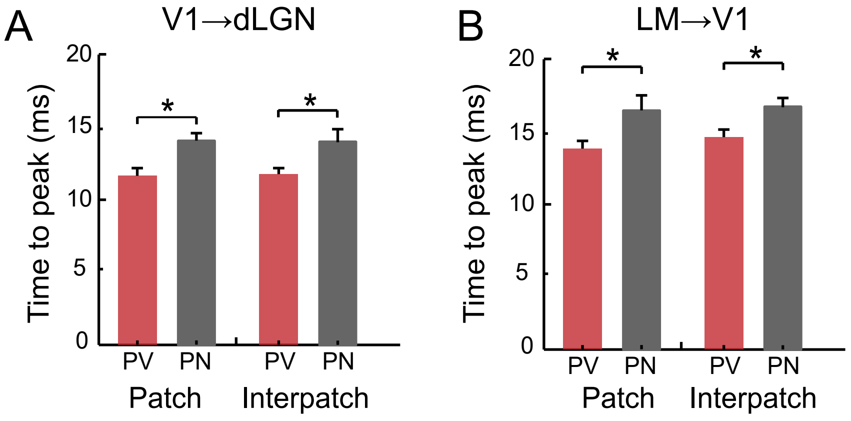

### Fig S6

Figure S6

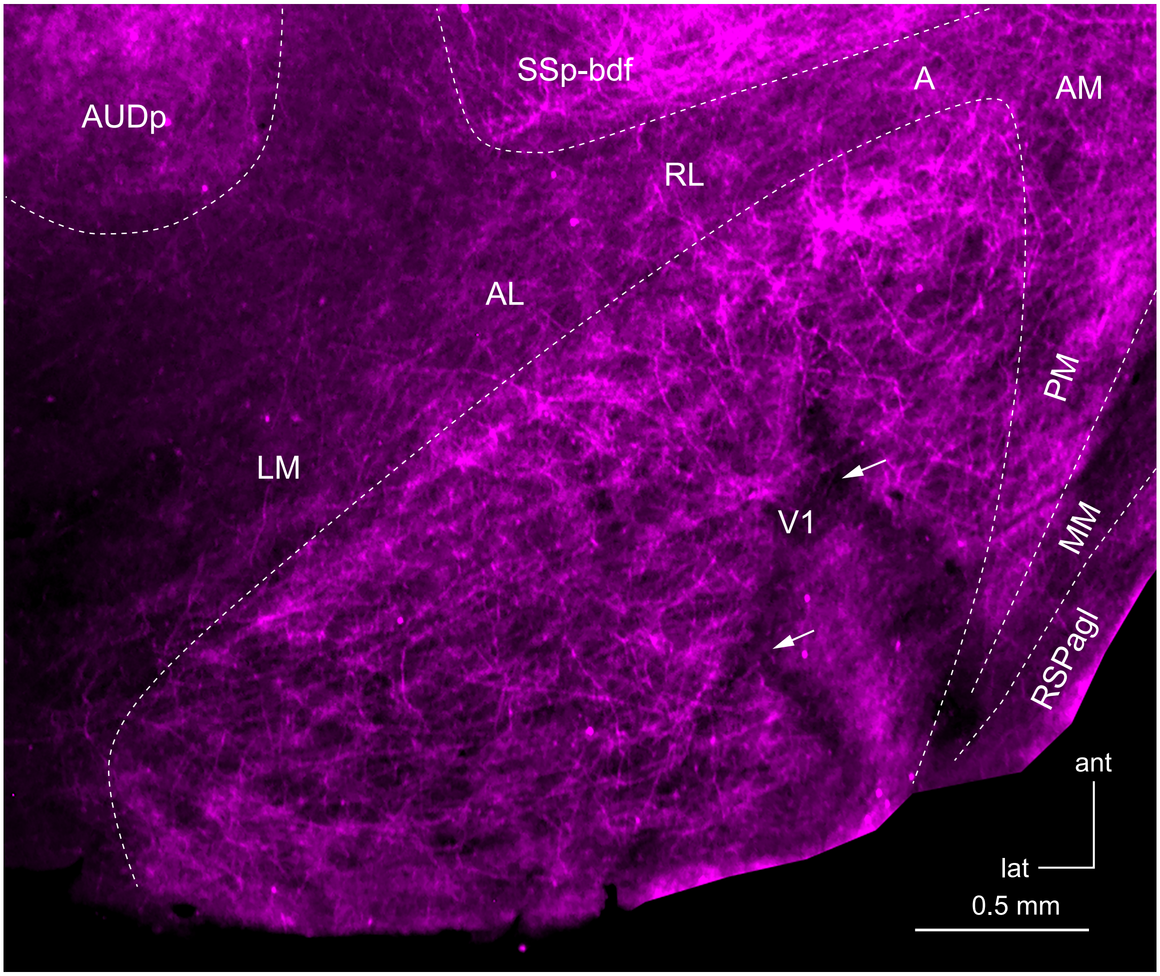

### Fig S7

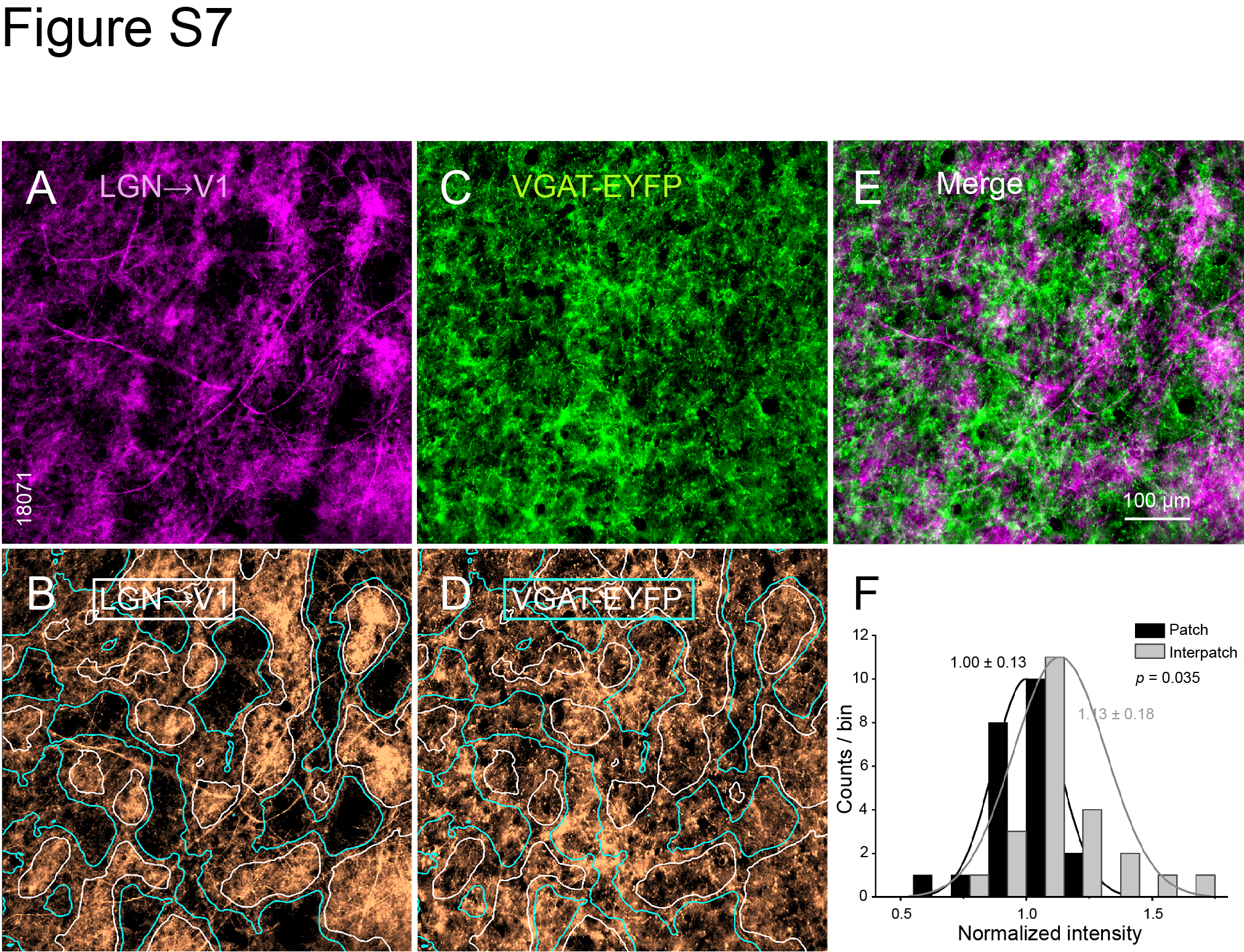

### Fig S8

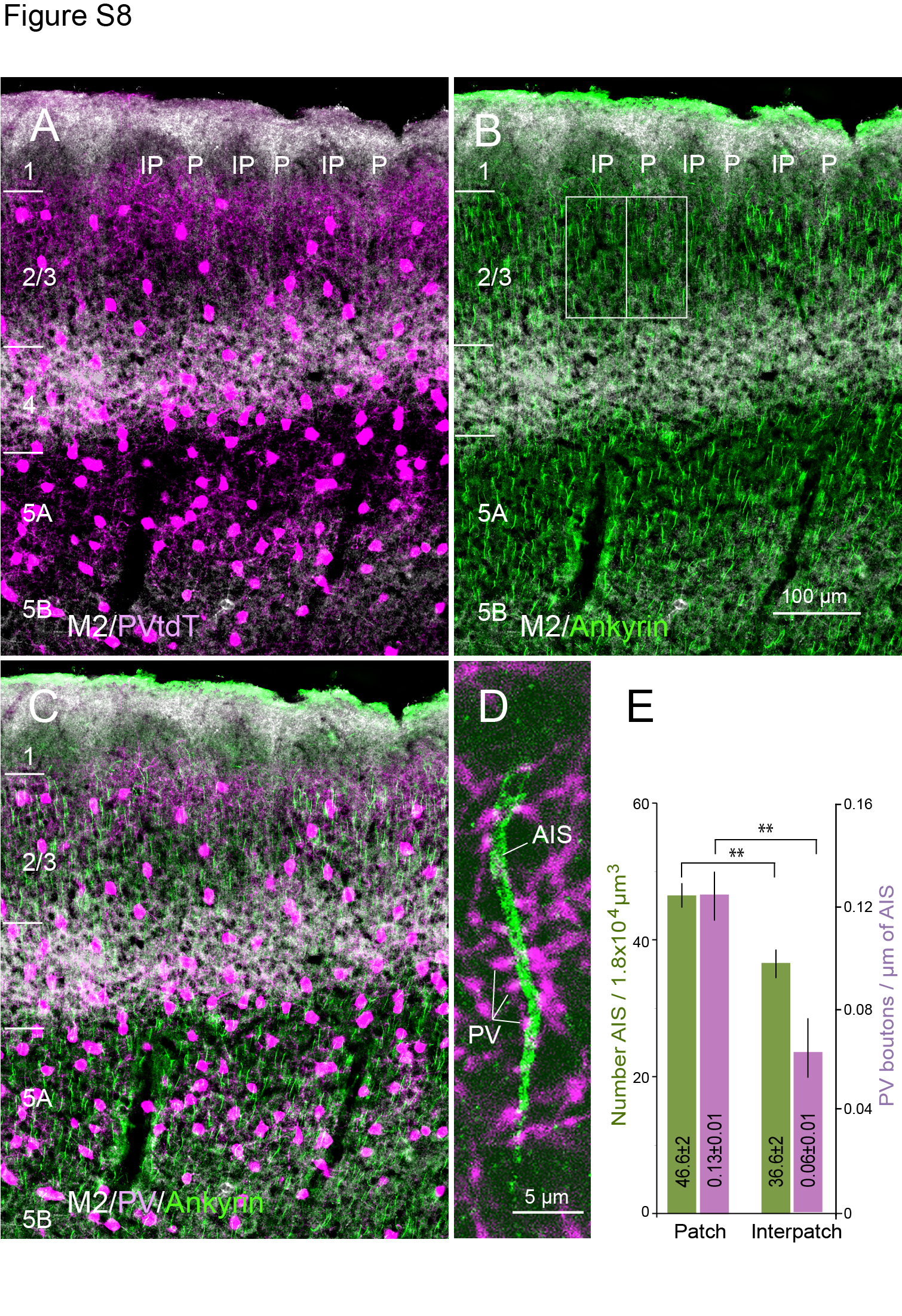
